# Hypoxia-related radiomics predict immunotherapy response: A multi-cohort study of NSCLC

**DOI:** 10.1101/2020.04.02.020859

**Authors:** Ilke Tunali, Yan Tan, Jhanelle E. Gray, Evangelia Katsoulakis, Steven A. Eschrich, James Saller, Theresa Boyle, Jin Qi, Albert Guvenis, Robert J. Gillies, Matthew B. Schabath

**Author notes:** Corresponding Author: Matthew B. Schabath, Ph.D., H. Lee Moffitt Cancer Center and Research Institute, 12902 Magnolia Drive MRC-CANCONT, Tampa, FL 33612.

## Abstract

Checkpoint blockade immunotherapy provides improved long-term survival in a subset of advanced stage non-small cell lung cancer (NSCLC) patients. However, highly predictive biomarkers of immunotherapy response are unmet clinical need. In this study, we utilized pre-treatment clinical factors and quantitative image-based biomarkers (radiomics) to identify a parsimonious model that predicts survival outcomes among NSCLC patients treated with immunotherapy. The NSCLC patients treated with single or double agent immunotherapy were included in three different cohorts: Training (N = 180), test (N = 90) and validation (N = 62) cohorts. The models were created based on overall survival (OS) and were additionally assessed for progression-free survival (PFS). Including most predictive radiomic features and clinical covariates, Classification and Regression Tree analysis was applied to stratify patients into survival risk-groups in the training cohort. The risk groups were later generated in the test and validation cohorts. Four independent NSCLC cohorts (total N = 446) were utilized for further validation of the radiomic signature. The biological underpinnings of the most informative radiomics were assessed using gene expression data from a radiogenomics dataset and validated by immunohistochemistry data (IHC). A parsimonious clinical-radiomics model was found to be significantly associated with OS and PFS after stratifying patients into groups of low-, moderate-, high-, and very-high risk of death and progression. This trained model was further tested and validated in two independent cohorts. When the extreme phenotypes were compared, the very-high risk group was found to be associated with extremely poor OS in both the test (hazard ratio [HR] = 5.35, 95% confidence interval [CI]: 2.14 – 13.36; 1-year OS = 11.1%) and validation (HR = 13.81, 95% CI: 2.58 – 73.93; 1-year OS = %47.6) cohorts when compared to the low risk group (HRs = 1.00; 1-year OS = 85.0% & 80.2%). Similar findings were observed for PFS. The final radiomic feature (GLCM inverse difference) was associated with OS in four independent NSCLC cohorts and was found to be positively associated with the hypoxia-related carbonic anhydrase, CAIX, by gene expression profiling and immunohistochemistry. We validated a novel clinical-radiomics model that is associated with OS and PFS among NSCLC patients treated with immunotherapy and identified a highly vulnerable subset of patients that are unlikely to respond to immunotherapy. The most informative radiomic feature was associated with CAIX, a marker of tumor hypoxia, tumor acidosis, and treatment resistance.

## Introduction

Checkpoint blockade immunotherapy has demonstrated durable clinical benefit in 20-50% of patients with advanced stage non-small-cell lung cancer (NSCLC)^1-6^. The patterns of immunotherapy response and progression are complex ^7^, including rapid disease progression^8^, hyperprogression^9^, and acquired resistance^10^. Because of this complexity in response patterns, there is a pressing challenge to identify robust predictive biomarkers that can identify patients that are more likely to benefit from immunotherapy. Tumor programmed cell death ligand-1 (PD-L1) expression by immunohistochemistry (IHC) is the only clinically approved biomarker to predict immunotherapy response however, recent clinical trials demonstrated significant improvements in clinical outcomes irrespective of PD-L1 expression level^6,11^. Furthermore, tumor mutational burden (TMB), defined as the total number of mutations per coding area of a tumor genome^12^, has shown to be a superior predictor of immunotherapy response in some studies^13-15^. Despite the potential clinical utility of TMB, there are limitations as tumor specimens have to be sufficient in both quantity and quality^14^. Additionally, tumors are evolutionarily dynamic and accumulate mutations rapidly^16^, and laboratory methods to calculate TMB can be timely and expensive which reduces the clinical utility of TMB. Moreover, tumor-based biomarkers, including both PD-L1 expression and TMB, are often subject to sampling bias due to molecular and cellular heterogeneity of the biopsied tumors^17^. As such, complimentary biomarkers that are predictive, non-invasive, and measured in a timely fashion would have direct translational implications. Quantitative image-based features, or radiomics^18^, reflect the underlying pathophysiology and tumor heterogeneity (Fig. 1) and have many advantages over tissue-based biomarkers as they can be rapidly calculated from standard-of-care medical images and reflect the entire region-of-interest (e.g., tumor) rather than the portion of the tumor that is assayed.

**Figure 1.**
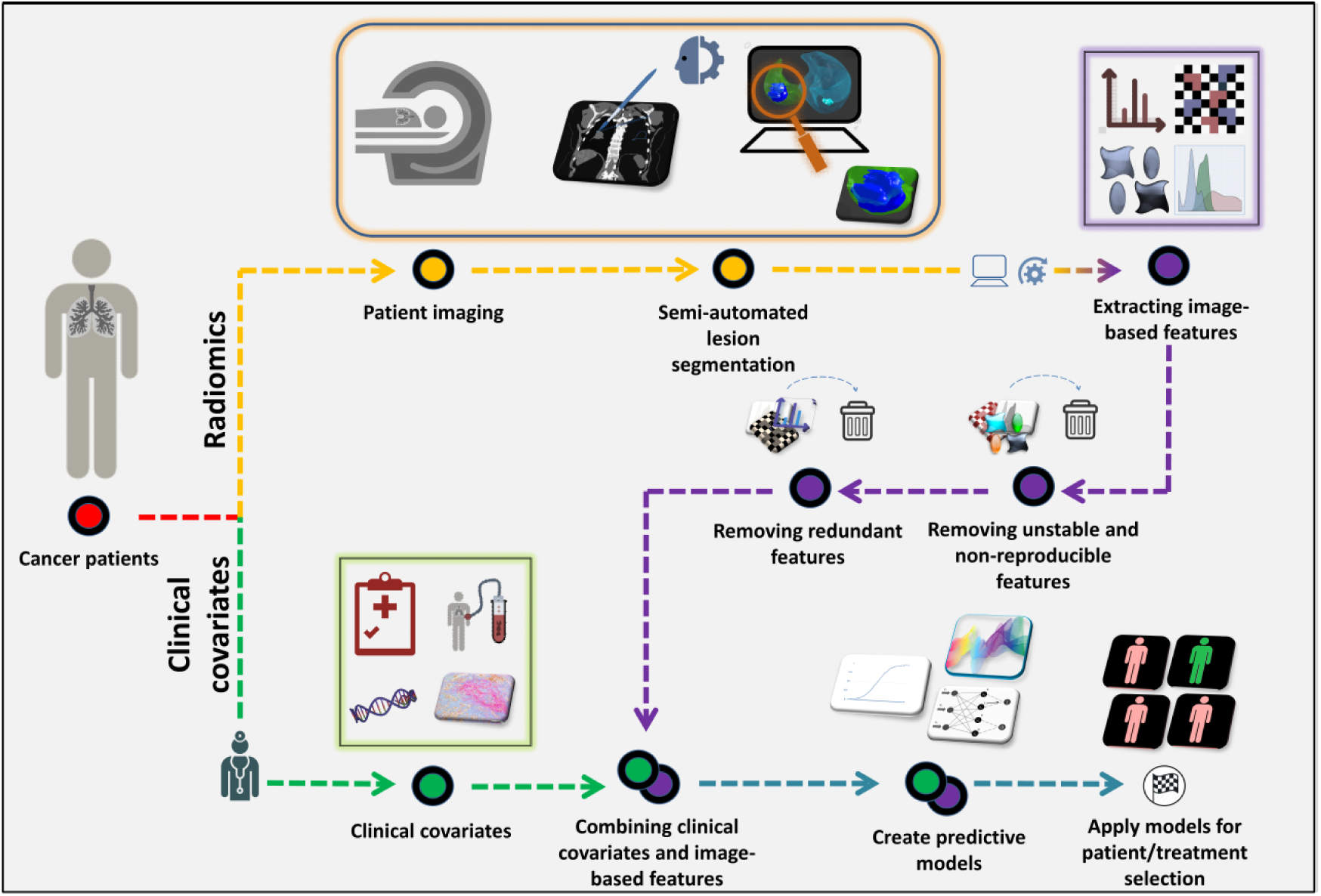
The Radiomics Pipeline. Pre-treatment (baseline) patient data are obtained, including: clinical covariates and computational image-based features (Radiomics). Radiomic features are extracted from standard-of-care imaging studies (yellow). Radiologists mark target lesions and lesions are automatically (or semi-automatically) segmented. Radiomic features are extracted from region-of-interest (purple). Unstable, non-reproducible and correlated radiomic features are removed. The remaining features are combined with the pre-treatment clinical covariates (green) and predictive model building approaches are applied which can be used for patient stratification and/or treatment selection.

In prior work^8^, we demonstrated the utility of radiomics of computed tomography (CT) images to predict rapid disease progression and hyperprogression phenotypes in NSCLC patients treated with immunotherapy. Building upon this work, in this present study we utilized pre-treatment clinical data and radiomic features extracted from CT scans to develop a parsimonious model that associates with survival outcomes among NSCLC patients treated with immunotherapy. The biological underpinnings of the radiomic features were assessed utilizing gene-expression information from a radiogenomics dataset and by IHC data. Furthermore, the radiomic features were assessed for survival in four independent NSCLC cohorts.

## Materials and Methods

### Immunotherapy-treated lung cancer patients

This analysis included 332 stage IIIB or IV NSCLC patients that were treated with immunotherapy using single agent programmed cell death-1 (Nivolumab, Pembrolizumab) or PD-L1 (Durvalumab, Atezolizumab), or combination of PD-L1 or programmed cell death-1 (PD-1) with cytotoxic T-lymphocyte-associated protein 4 (Ipilimumab, Tremelimumab) as second agent. Inclusion criteria included patients having a baseline CT or PET/CT scan prior to the initiation of checkpoint blockade immunotherapy in less than 90 days and at least one measurable lung lesion defined by Response Evaluation Criteria in Solid Tumors (RECIST) criteria. The patients were divided into training (MCC 1, N = 180), test (MCC 2, N = 90) and validation cohorts (VA, N = 62). Patients in the training cohort were enrolled in clinical trials treated between June 2011 and January 2016 at Moffitt Cancer Center. Details of the training cohort have been previously published^8^. Patients in the test cohort were treated with immunotherapy between May 2015 and October 2017 where 94.6% were treated as standard-of-care and 5.4% were enrolled in clinical trials at Moffitt Cancer Center. Patients in the validation cohort were treated with standard-of-care immunotherapy between July 2015 and February 2019 at James A. Haley Veterans’ Hospital.

Patient data were obtained from electronic medical records and institutional databases and included demographics, stage of diseases, histology, treatment, vital status, targeted mutations, ECOG performance status, RECIST, hematology data, vital status (deceased or alive), and date of death or last follow-up. Dates of progressions were abstracted and defined as progressive disease using RECIST or iRECIST definition or clinical progression evaluated by the treating clinician whenever RECIST or iRECIST data were not available. Details of data elements are provided in the Supplementary Methods. This study was approved with a waiver of informed consent by the University of South Florida Institutional Review Board.

### Radiogenomics dataset

A previously described dataset^19^ of 103 surgically resected adenocarcinoma patients who had pre-surgical CT scans and gene-expression data were used to identify potential biological underpinnings of most informative radiomic features. Briefly, gene expression was IRON-normalized and batch-corrected for RNA quality Pathway and Gene Ontology Enrichment was performed using Clarivate Analytics MetaCore^19^.

### Immunohistochemistry

To validate the findings from the radiogenomics dataset, 16 NSCLC patients who had surgically resected tumors and pre-surgical CT scans were identified for IHC staining. Details of the staining procedure are provided in the Supplemental Methods. The stainings were quantified by two approaches: i) an automated evaluation of positive staining percentage defined by the Aperio ImageScope^20^ Positive Pixel Count algorithm as the total number of positive stained pixels by total number of tumor and its immediate microenvironment pixels; and ii) the H-scoring metric^21^ quantified by a board-certified pathologist (J.S.) who was blinded to radiomics and Aperio ImageScope scoring data.

### Prognostic validation datasets

The radiomics data were further validated in four published NSCLC non-immunotherapy treated patient datasets. Only OS was available for these datasets. The first dataset^22,23^ included 62 adenocarcinoma patients who underwent surgical resection as first course therapy at Moffitt Cancer Center and had pre-surgical CT scans within 2 months prior to surgery. The second dataset^22,23^ included 47 adenocarcinoma patients who underwent surgical resection as first course therapy at the Maastricht Radiation Oncology Clinic and had pre-surgical CT scans within 2 months prior to surgery. The third dataset included 234 patients^24,25^ diagnosed with screen-detected incident lung cancers in the National Lung Screening Trial. The fourth dataset included 103 adenocarcinoma patients from the radiogenomics dataset that was described above.

### Tumor segmentation and radiomics extraction

Figure 1 presents an overview of the radiomics pipeline used in this study. Pre-treatment contrast-enhanced thoracic CT scans performed ≤ 90 days prior to the initiation of immunotherapy (baseline) were retrieved from the picture archiving and communication system and loaded into HealthMyne^®^ Quantitative Imaging Decision Support (QIDS) software (https://www.healthmyne.com). Two radiologists (Y.T. and J.Q.) with more than 10 years of clinical experience selected the largest lung tumors of each patient and initialized an automated 3D segmentation algorithm using the HealthMyne^®^ QIDS Rapid Precise Metrics software. The tumor delineation output of the 3D segmentation algorithm was either confirmed or edited whenever necessary by a radiologist.

The tumor mask images (i.e., tumor delineations) were imported into in-house radiomic feature extraction toolbox that was created using MATLAB^®^ 2015b (The Mathworks Inc., Natick, Massachusetts) and C++ (https://isocpp.org). To standardize spacing across all images, all CT images were resampled to a single voxel spacing of 1mm x 1mm x 1mm using cubic interpolation. For texture features, Hounsfield units (HUs) were resampled into fixed bin sizes of 25 HUs discretized from –1000 to 1000 HU.

A total of 213 radiomic features were extracted from intratumoral region (N = 122 features) and the peritumoral region 3 mm outside of tumor boundary (N = 91 features) using standardized algorithms from the Image Biomarker Standardization Initiative (IBSI) v5^26^. Peritumoral regions were bounded by lung parenchyma masks to exclude any peritumoral region that exceed outside of lung parenchyma. Unstable and non-reproducible radiomic features were eliminated using methods described elsewhere^27^ utilizing two publicly available datasets^28,29^. Details on feature selection are mentioned in Supplementary methods.

### Statistical analysis

All statistical analyses were performed using Stata/MP 14.2 (StataCorp LP, College Station, Texas) and R Project for Statistical Computing version 3.4.3 (http://www.r-project.org/). Differences for clinical covariates were tested using Fisher’s exact test for categorical variables and Mann-Whitney *U* test and Student’s *t*-test for continuous variables. Survival analyses were performed using Kaplan-Meier survival estimates and the log-rank tests. The OS and progression-free survival (PFS) were two dependent variables. For OS, an event was defined as death and the data were right censored at 36-months. For PFS, an event was defined as death, clinical progression or RECIST based progression of cancer and the data were right-censored at 36 months. The index date for OS and PFS was the initiation of immunotherapy.

A rigorous model building approach was employed to reduce the number of covariates and identify the most prognostic clinical covariates and radiomic features. For the clinical covariates, univariable Cox regression was performed and covariates significantly (P < 0.05) associated with OS were retained. To produce a parsimonious clinical model, remaining clinical covariates were included in a stepwise backward elimination Cox regression model using a threshold of 0.01 for inclusion. For the radiomic features, univariable Cox regression was performed and radiomic features significantly associated with OS after Bonferroni-Holm correction (P < 0.05) were retained. Radiomic features correlated with tumor volume (Pearson’s correlation coefficient ≥ 0.80) were removed. Among the remaining radiomic features, correlated features were identified using absolute Pearson’s correlation coefficient ≥ 0.80 and the feature with the smallest P-value from the univariable analysis was retained. The remaining radiomic features, were utilized to identify a parsimonious radiomics model using a stepwise backward elimination approach applying a threshold of 0.01 for inclusion. The final covariates from the clinical model and the final features from the radiomics model were combined and Classification and Regression Tree (CART) was used to stratify patients into risk groups. CART is a non-parametric approach modified for failure time data^30^ that classifies variables through a decision tree composed of splits or nodes, where the split points are optimized based on impurity criterion. The clinical-radiomics CART model found using training cohort (MCC 1) was tested and validated utilizing test (MCC 2) and validation (VA) cohorts. Time-dependent AUCs and confidence intervals (CIs) were calculated for 6, 12, 24 and 36 months for training and test cohorts. The most predictive radiomics feature was also validated in four independent cohorts.

For the radiogenomics analysis, the highest prognostic radiomic feature was compared to every gene probesets using two different approaches: correlation and two-group analysis. For the correlation analysis, gene probesets were filtered and determined as significant using the following criteria: Pearson’s correlation with a threshold |R|>0.4, an expression filter with max expression of gene > 5, and an inter-quartile filter (interquartile range > log2 (1.2 fold-changes). Gene probesets were filtered and determined as significant using the following criteria based on a Student’s t test p < 0.001 and mean log fold-changes (LFC) between high and low prognostic radiomic feature of LFC > log2 (1.4 fold-changes). The significant probesets from the two analyses were intersected yielding a final list of probesets significantly associated with the most prognostic radiomic feature.

### Radiomics Quality Score

Radiomics Quality Score (RQS) was calculated using the established metric developed by Lambin et al. ^31^ which assesses the quality of a radiomics study in order to minimize bias and enhance the usefulness of the radiomics models. The RQS contains 16 key components where a maximum score of 36 could be achieved.

## Results

### *Immunotherapy treated patient demographics* (MCC 1, MCC 2, and VA)

Type of checkpoint inhibitor, ECOG performance status, number of previous lines of therapy, serum albumin, lymphocyte counts, and neutrophils to lymphocytes ratio (NLR) were significantly different between the training (MCC 1) and test cohorts (MCC 2, Table 1). Also, significant differences were found for OS and PFS between training and test cohorts (36-month OS 32.6% vs. 19.2%, respectively; 36-month PFS 20.8% vs. 9.5%, respectively; Table 2 & Supplementary Fig. 1).

**Table 1.** Patient characteristics by the training, test and validation cohorts.

| Characteristic | Training Cohort<br>(MCC 1)<br>(N = 180) | Test Cohort<br>(MCC 2)<br>(N = 90) | P-Value<sup>1</sup> | Validation<br>cohort (VA)<br>(N = 62) | P-Value<sup>2</sup> |
| --- | --- | --- | --- | --- | --- |
| **Age at initiation of treatment, N (%)** |  |  |  |  |  |
| *Dichotomized* |  |  |  |  |  |
| < 65 | 68 (37.8) | 37 (41.1) |  | 15 (24.2) |  |
| ≥ 65 | 112 (62.2) | 53 (58.9) | 0.599 | 47 (75.8) | 0.063 |
| **Median, (95% CI)** | 67 (65-68) | 67 (64-69) | 0.783 | 68 (67-71) | **0.026** |
| **Sex, N (%)** |  |  |  |  |  |
| Female | 95 (52.8) | 43 (47.8) |  | 3 (4.8) |  |
| Male | 85 (47.2) | 47 (52.2) | 0.442 | 59 (95.2) | **<0.001** |
| **Smoking status<sup>1</sup>, N (%)** |  |  |  |  |  |
| Never smoker | 30 (16.7) | 16 (17.8) |  | 2 (3.2) |  |
| Ever smoker | 146 (81.1) | 74 (82.2) | 0.866 | 60 (96.8) | **0.004** |
| Unknown/Missing | 4 (2.2) | 0 (0) |  | 0 (0) |  |
| **Stage, N (%)** |  |  |  |  |  |
| III | 6 (3.3) | 4 (4.4) |  | 13 (21.0) |  |
| IV | 174 (96.7) | 86 (95.6) | 0.735 | 49 (79.0) | **<0.001** |
| **Histology, N (%)** |  |  |  |  |  |
| Adenocarcinoma/others | 137 (76.1) | 71 (78.9) |  | 43 (69.3) |  |
| Squamous cell carcinoma | 43 (23.9) | 19 (21.1) | 0.648 | 19 (30.7) | 0.135 |
| **Checkpoint inhibitors, N (%)** |  |  |  |  |  |
| Anti PD-L1 | 48 (26.6) | 18 (20.0) |  | 8 (12.9) |  |
| Anti PD-1 | 57 (31.7) | 69 (76.7) |  | 54 (87.1) |  |
| Doublet | 75 (41.7) | 3 (3.3) | **<0.001** | 0 (0) | **<0.001** |
| **ECOG performance status, N (%)** |  |  |  |  |  |
| 0 | 39 (21.7) | 10 (11.1) |  | 12 (19.4) |  |
| 1 | 141 (78.3) | 67 (74.4) |  | 39 (62.9) |  |
| 2 | 0 (0) | 13 (14.4) | **<0.001** | 11 (17.7) | **<0.001** |
| **Previous lines of therapy on current diagnosis, N (%)** |  |  |  |  |  |
| None | 70 (43.9) | 21 (23.3) |  | n/a |  |
| 1 | 48 (26.7) | 47 (52.2) |  | n/a |  |
| ≥ 2 | 62 (34.4) | 22 (24.4) | **<0.001** | n/a | - |
| **Number of metastatic sites, N (%)** |  |  |  |  |  |
| 1 | 82 (46.6) | 51 (56.7) |  | 25 (40.3) |  |
| ≥ 2 | 98 (54.4) | 39 (43.3) | 0.094 | 37 (59.7) | 0.554 |
| **EGFR mutational status<sup>3</sup>, N (%)** |  |  |  |  |  |
| Not Detected | 107 (59.4) | 37 (41.1) |  | n/a |  |
| Detected | 25 (13.9) | 5 (5.6) |  | n/a |  |
| Missing/Inconclusive | 48 (26.7) | 48 (53.3) | 0.355 | n/a | - |
| **KRAS mutational status<sup>3</sup>, N (%)** |  |  |  |  |  |
| Not Detected | 61 (33.9) | 20 (22.2) |  | n/a |  |
| Detected | 29 (16.1) | 12 (13.3) | 0.664 | n/a |  |
| Missing/Inconclusive | 90 (50.0) | 58 (64.4) |  | n/a | - |
| **Hematology, median, (95% CI)** |  |  |  |  |  |
| Serum albumin, (g/dL) | 4.0 (3.9-4.0) | 3.8 (3.6-3.9) | **<0.001** | 3.9 (3.7-4.0) | 0.087 |
| Lymphocytes, (1e+9/L) | 1.3 (1.2-1.4) | 1.0 (0.9-1.2) | **<0.001** | 1.0 (0.9-1.2) | **0.014** |
| WBC, (1e+9/L) | 7.1 (6.7-7.6) | 7.7 (6.8-8.8) | 0.246 | 7.5 (6.7-8.7) | 0.383 |
| Neutrophils, (1e+9/L) | 4.8 (4.4-5.1) | 5.3 (4.6-6.5) | 0.131 | 5.6 (4.8-6.1) | 0.329 |
| Ratio of: Neutrophils/Lymphocytes | 3.7 (3.2-4.1) | 5.2 (4.0-7.5) | **0.002** | 5.3 (4.1-6.8) | **0.004** |

Abbreviations: CI = confidence interval;
**Bold** P-values are statistically significant; P-values for continuous variables were calculated using Mann-Whitney test and Fisher's Exact Test for categorical variables.
<sup>1</sup>P-values were calculated comparing training cohort and the test cohorts.
<sup>2</sup>P-values were calculated comparing training cohort and the validation cohorts.
<sup>3</sup>P-values for smoking status, *EGFR* mutational status and *KRAS* mutational status were calculated for patients without missing/inconclusive data.
Majority of the training (95.3%) and the test cohorts (86.7%) were self-reported White race and majority of the training (97.0%) and the test cohorts (88.9%) were self-reported non-Hispanic. For the validation cohort the racial and ethnicity data were not available.

**Table 2.** Overall survival and progression free survival rates by training and test cohorts and patient risk groups^1^

|  | Percent survival at: |  |  |  |
| --- | --- | --- | --- | --- |
|  | 6 months | 12 months | 24 months | 36 months |
| **<u>Overall survival</u>** |  |  |  |  |
| ***Overall by cohort*** |  |  |  |  |
| Training cohort (MCC 1) | 82.7% | 60.8% | 42.1% | 32.6% |
| Test cohort (MCC 2) | 61.7% | 46.2% | 22.4% | 19.2% |
| Validation cohort (VA) | 75.5% | 63.0% | 31.9% | 12.0% |
| ***By risk group*** |  |  |  |  |
| **Low-risk** |  |  |  |  |
| Training cohort (MCC 1) | 100% | 95.2% | 84.7% | 84.7% |
| Test cohort (MCC 2) | 95.0% | 85.0% | 38.9% | 38.9% |
| Validation cohort (VA) | 91.7% | 80.2% | 80.2% | 40.1% |
| **Moderate-risk** |  |  |  |  |
| Training cohort (MCC 1) | 92.6% | 76.4% | 59.7% | 47.9% |
| Test cohort (MCC 2) | 67.8% | 56.7% | 33.1% | n/a |
| Validation cohort (VA) | 72.2% | 66.2% | 29.8% | n/a |
| **High-risk** |  |  |  |  |
| Training cohort (MCC 1) | 81.1% | 54.4% | 24.9% | 15.6% |
| Test cohort (MCC 2) | 62.1% | 34.1% | 17.1% | 8.5% |
| Validation cohort (VA) | 74.4% | 57.2% | 32.7% | n/a |
| **Very-high-risk** |  |  |  |  |
| Training cohort (MCC 1) | 59.9% | 24.3% | 12.2% | 0% |
| Test cohort (MCC 2) | 16.7% | 11.1% | 0% | 0% |
| Validation cohort (VA) | 66.7% | 47.6% | n/a | n/a |
| **<u>Progression-free survival</u>** |  |  |  |  |
| ***Overall by cohort*** |  |  |  |  |
| Training cohort (MCC 1) | 47.9% | 32.8% | 22.8% | 20.8% |
| Test cohort (MCC 2) | 37.9% | 19.6% | 9.5% | 9.5% |
| ***By risk group*** |  |  |  |  |
| **Low-risk** |  |  |  |  |
| Training cohort (MCC 1) | 71.4% | 71.4% | 65.5% | 65.5% |
| Test cohort (MCC 2) | 73.7% | 46.3% | 29.8% | 29.8% |
| **Moderate-risk** |  |  |  |  |
| Training cohort (MCC 1) | 65.9% | 43.0% | 31.2% | 25.0% |
| Test cohort (MCC 2) | 23.9% | 19.1% | 9.6% | n/a |
| **High-risk** |  |  |  |  |
| Training cohort (MCC 1) | 43.0% | 27.0% | 9.8% | 9.8% |
| Test cohort (MCC 2) | 41.3% | 15.0% | 3.8% | n/a |
| **Very-high-risk** |  |  |  |  |
| Training cohort (MCC 1) | 15.7% | 0% | 0% | 0% |
| Test cohort (MCC 2) | 11.1% | 0% | 0% | 0% |

<sup>1</sup>Cells were marked as n/a whenever all of the patients were censored for the given interval.

**Table 3.**
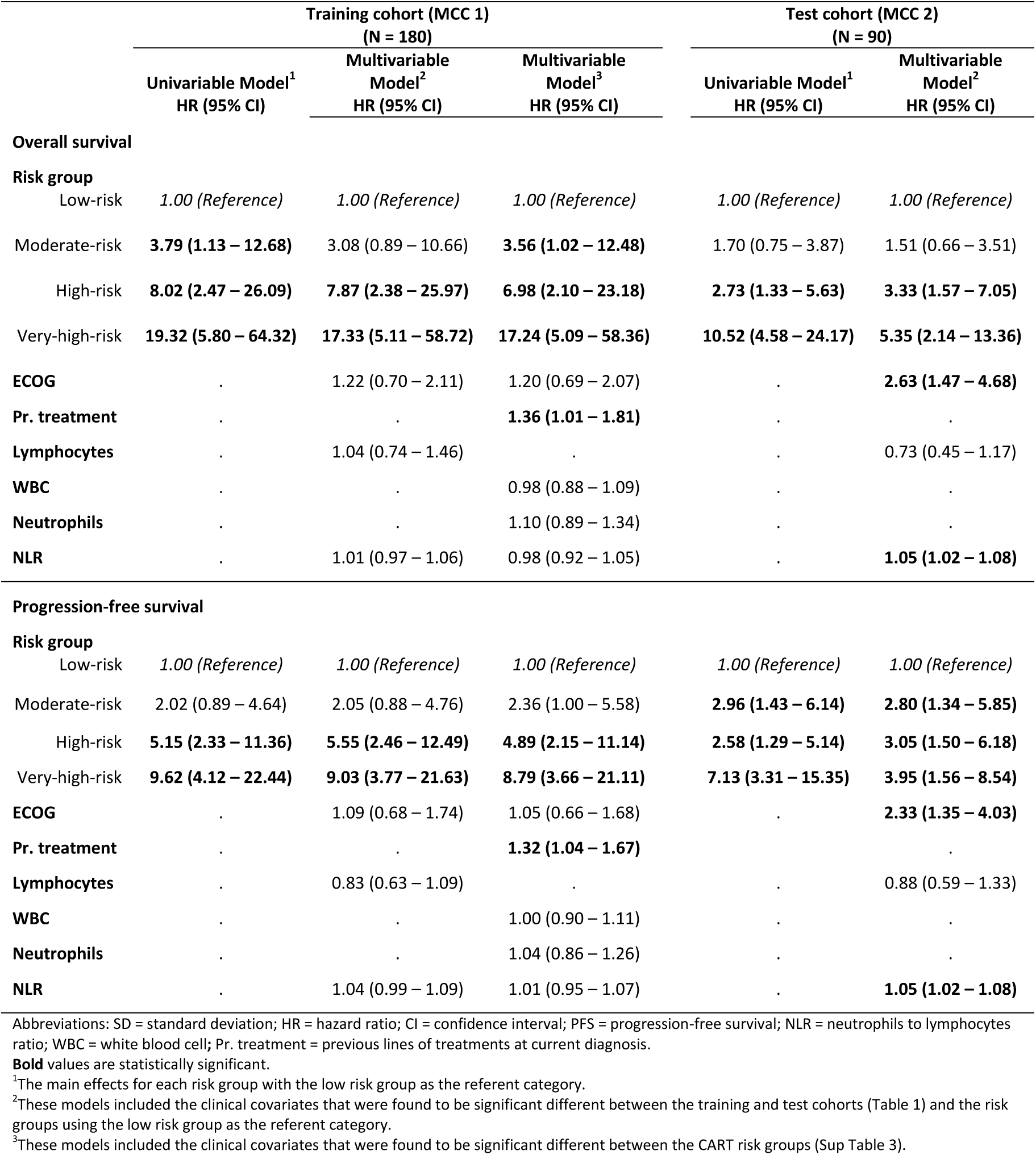
Univariable and multivariable Cox regression analysis for overall survival and progression-free survival for the training and test cohorts.

Median age, sex, smoking status, stage, type of checkpoint inhibitor, ECOG performance status, lymphocyte counts, and NLR were significantly different between the training (MCC 1) and validation (VA) cohorts (Table 1). However, OS was not significantly different between the two cohorts (Supplementary Fig. 1).

### Clinical model

Among the 16 clinical covariates that were considered for the clinical model (Table 1), four clinical features (serum albumin, number of metastatic sites, previous lines of therapy and neutrophils counts) were significantly associated with OS in univariable analysis utilizing the training cohort (MCC 1). After utilizing a stepwise backward elimination model, the final parsimonious clinical model included two clinical features: serum albumin (hazard ratio [HR] = 0.33; 95% CI: 0.20-0.52) and number of metastatic sites (HR = 2.14; 95% CI: 1.48-3.11).

### Radiomics model

Among 213 intratumoral and peritumoral radiomic features, 67 features were found to be stable and reproducible (Fig. 2a). Eight of the 67 features were removed because they were correlated with tumor volume. In univariable analysis, twelve features were found to be significantly associated with OS and ten of these features were removed because of high correlation (Fig. 2b). Among the two features remaining (gray level co-occurrence matrix [GLCM] inverse difference and peritumoral quartile coefficient), GLCM inverse difference was found as the most informative radiomic feature (HR = 1.41; 95% CI: 1.19-1.67, p < 0.001) after a stepwise backward elimination.

**Figure 2A).**
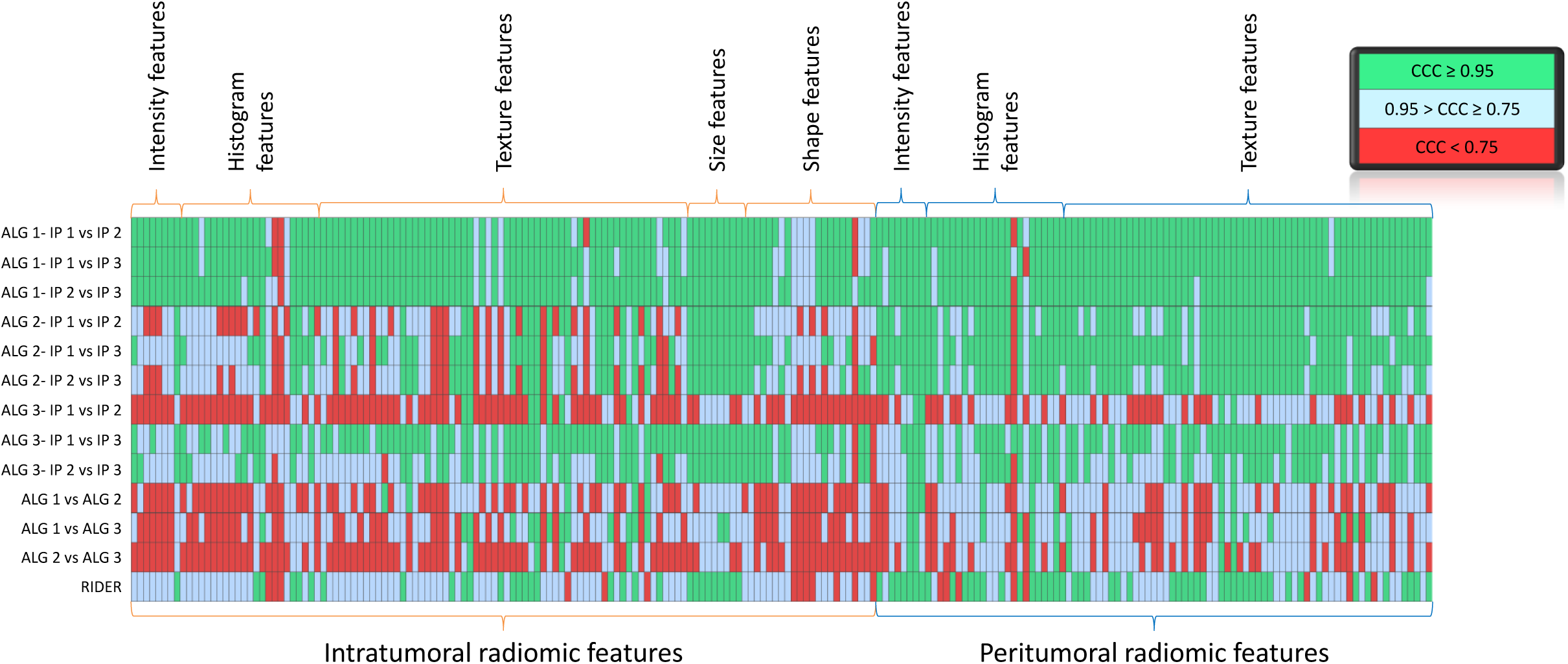
Heat map of concordance correlation coefficients for different segmentations and image acquisitions of radiomic features. Each column in the heat map represents a radiomic feature from the indicated feature group and region-of-interest (e.g., intratumoral or peritumoral). The features are compared between different segmentation algorithms (ALG), different initial parameters (IP) and test-retest scans (RIDER). The green boxes represent higher (CCC > 0.95), blue boxes represent moderate (CCC ≥ 0.75 & CCC ≤ 0.95) and red boxes represent lower (CCC < 0.75) CCCs.

**Figure 2B).**
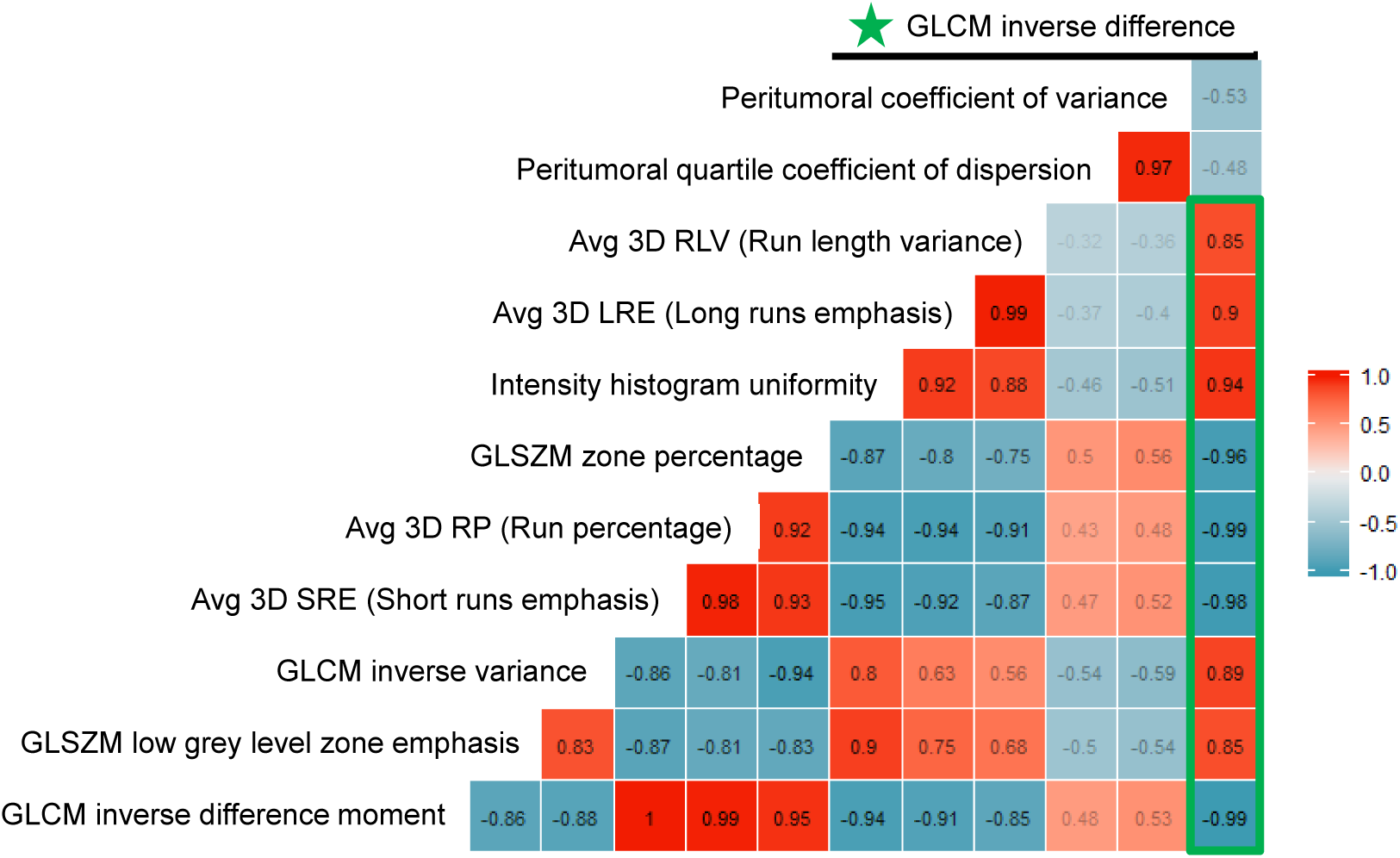
Correlation matrix for the radiomic features that were significantly associated with overall survival in the univariable analysis. The feature in the final parsimonious model GLCM inverse difference and it is found to be correlated with nine other features shown inside the green box.

### CART analysis

Based on most informative two clinical covariates and one radiomic feature, CART analysis have found novel cut points (Fig. 2c) and grouped patients into six risk groups (Supplementary Fig. 2). These groups were further collapsed into four risk groups: low risk, moderate risk, high risk, and very-high risk (Fig 3a). The risk groups identified in the training cohort (MCC 1) were also extracted for test (MCC 2) and validation (VA) cohorts (Table 2 and Fig. 3b&c) and time-dependent AUCs were calculated for OS (Supplementary Fig. 3). Specifically,, the model achieved an AUC of 0.784 (95% CI: 0.693 – 0.876) for 6 months OS and an AUC of 0.716 (95% CI: 0.558 – 0.843) for 24 months utilizing the test cohort. When PFS was used, similar findings were observed for the training (MCC 1) and the test (MCC 2) cohorts (Fig 3d&e). We did not have PFS data for the validation cohort (VA).

**Figure 2C).**
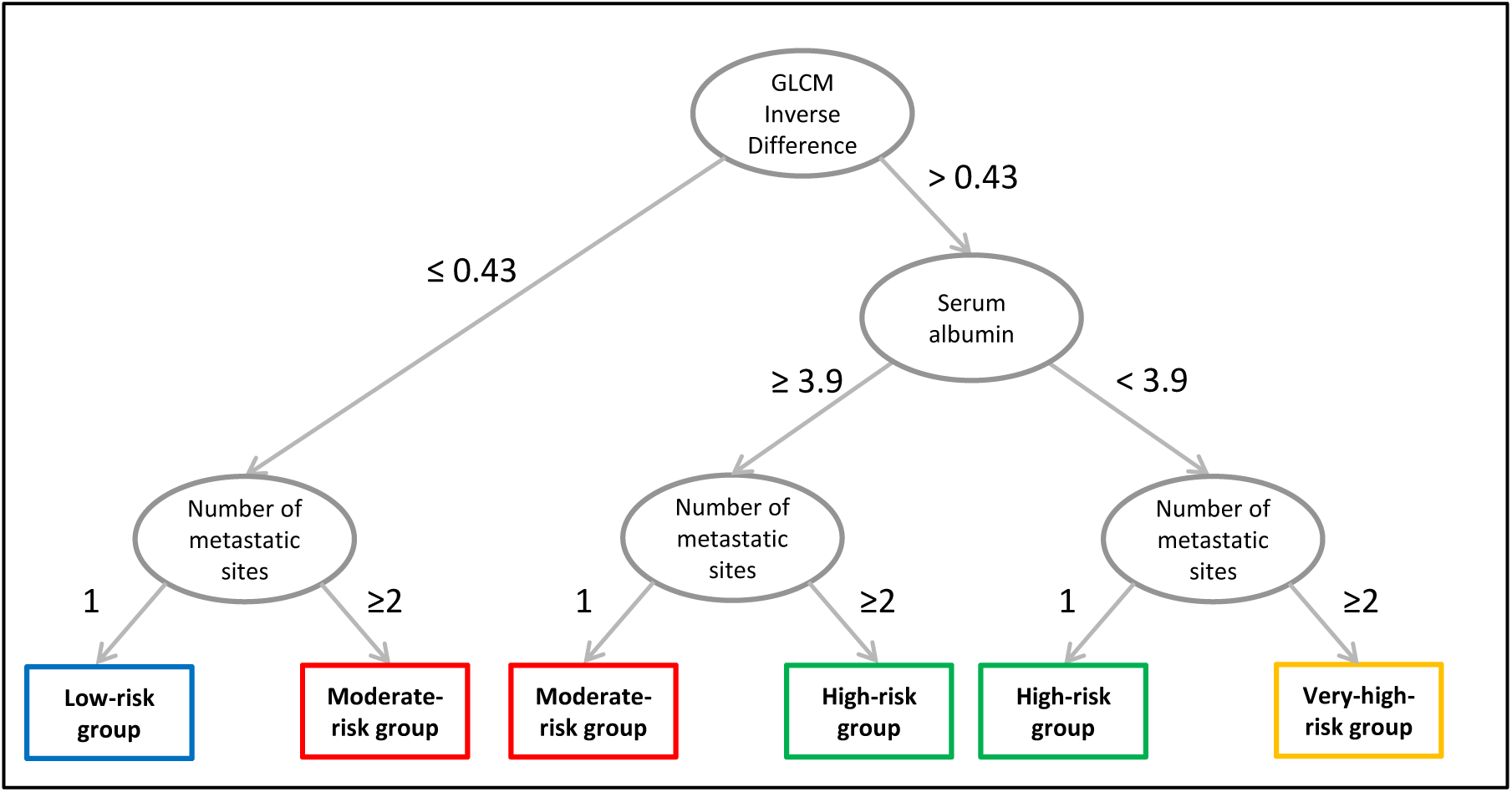
The Classification and Regression Tree (CART) was used to identify patient risk groups based on a model containing one radiomic feature and two clinical features. Patients were grouped from low risk to very high risk based on the CART decision nodes and terminal nodes.

**Figure 3.**
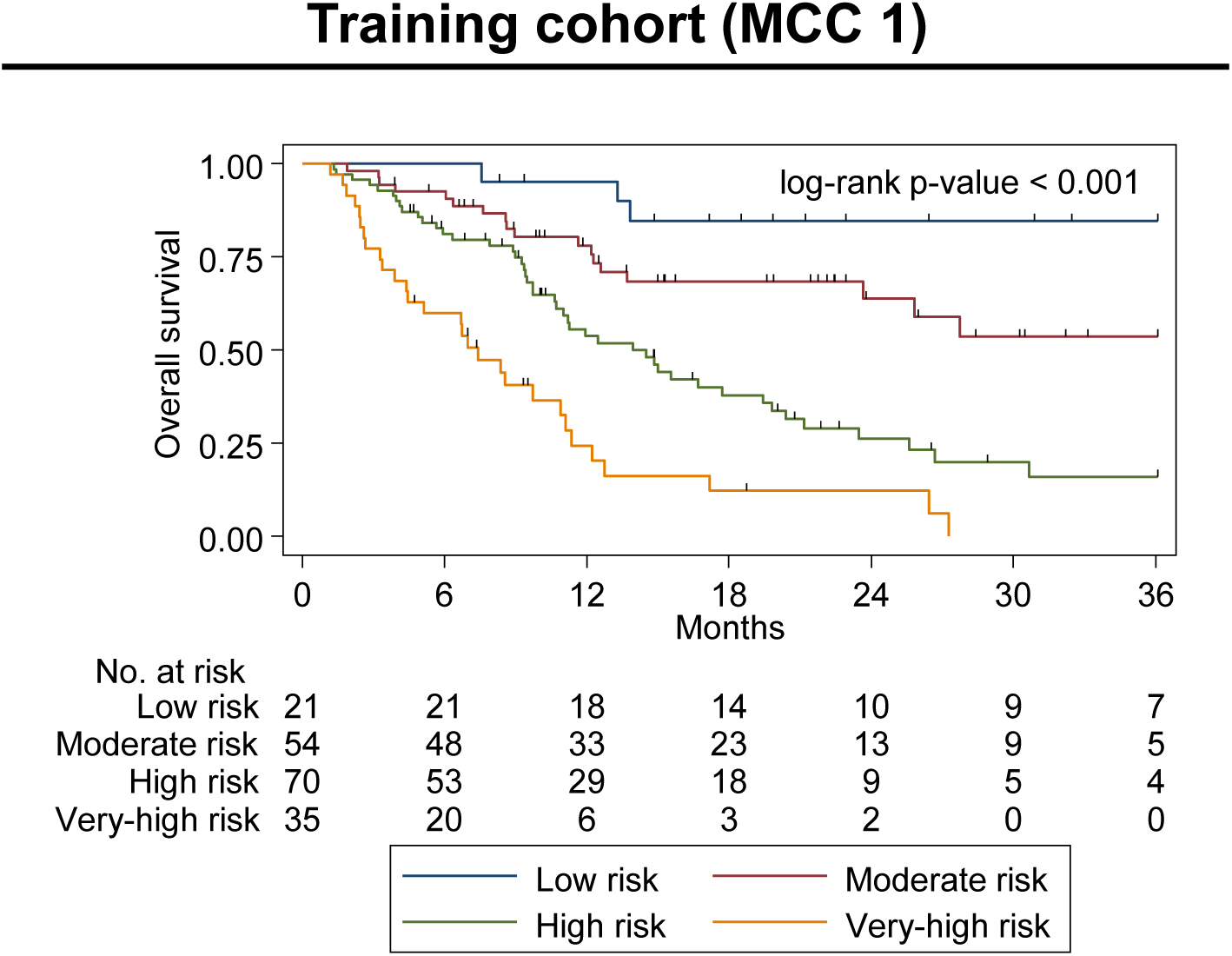
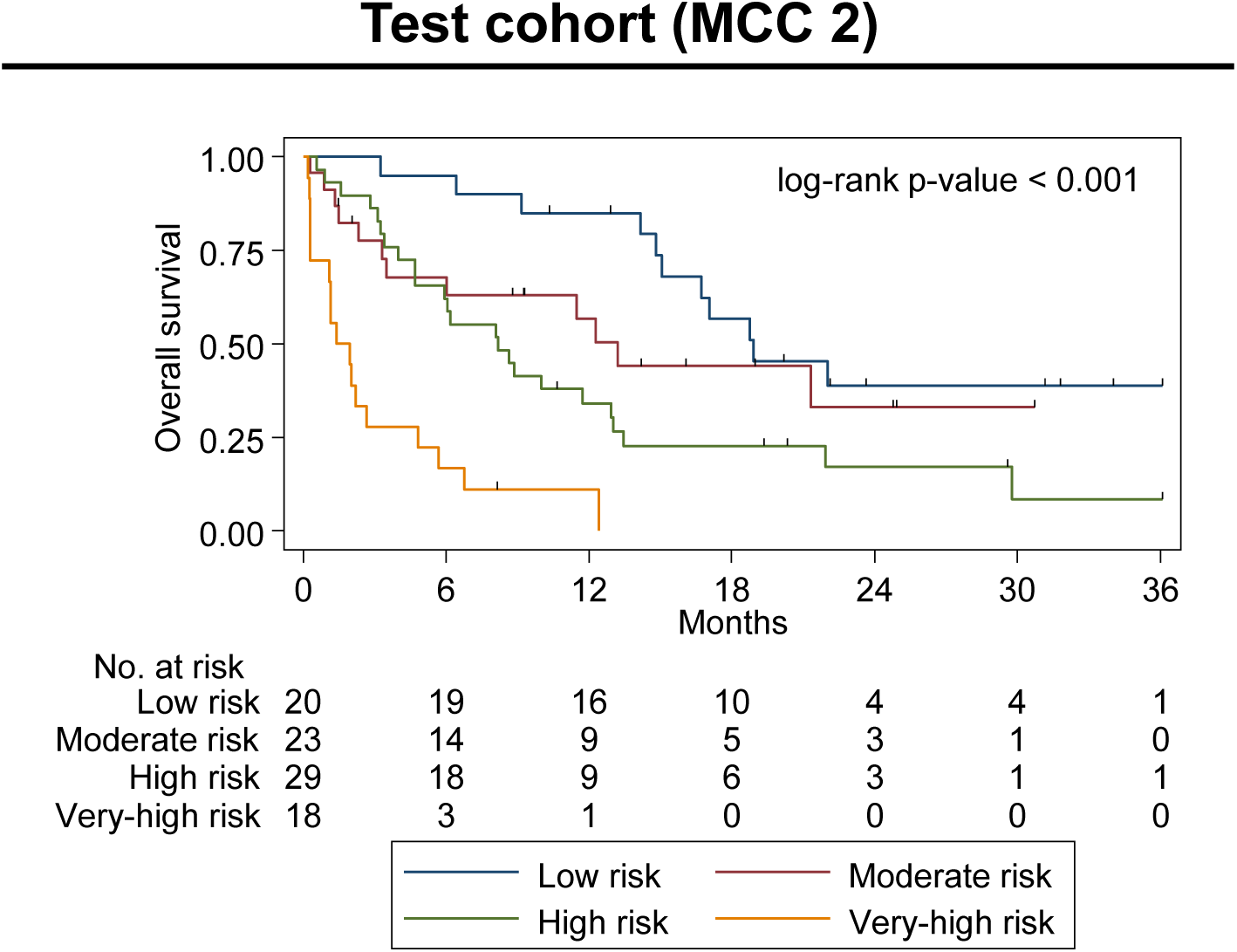
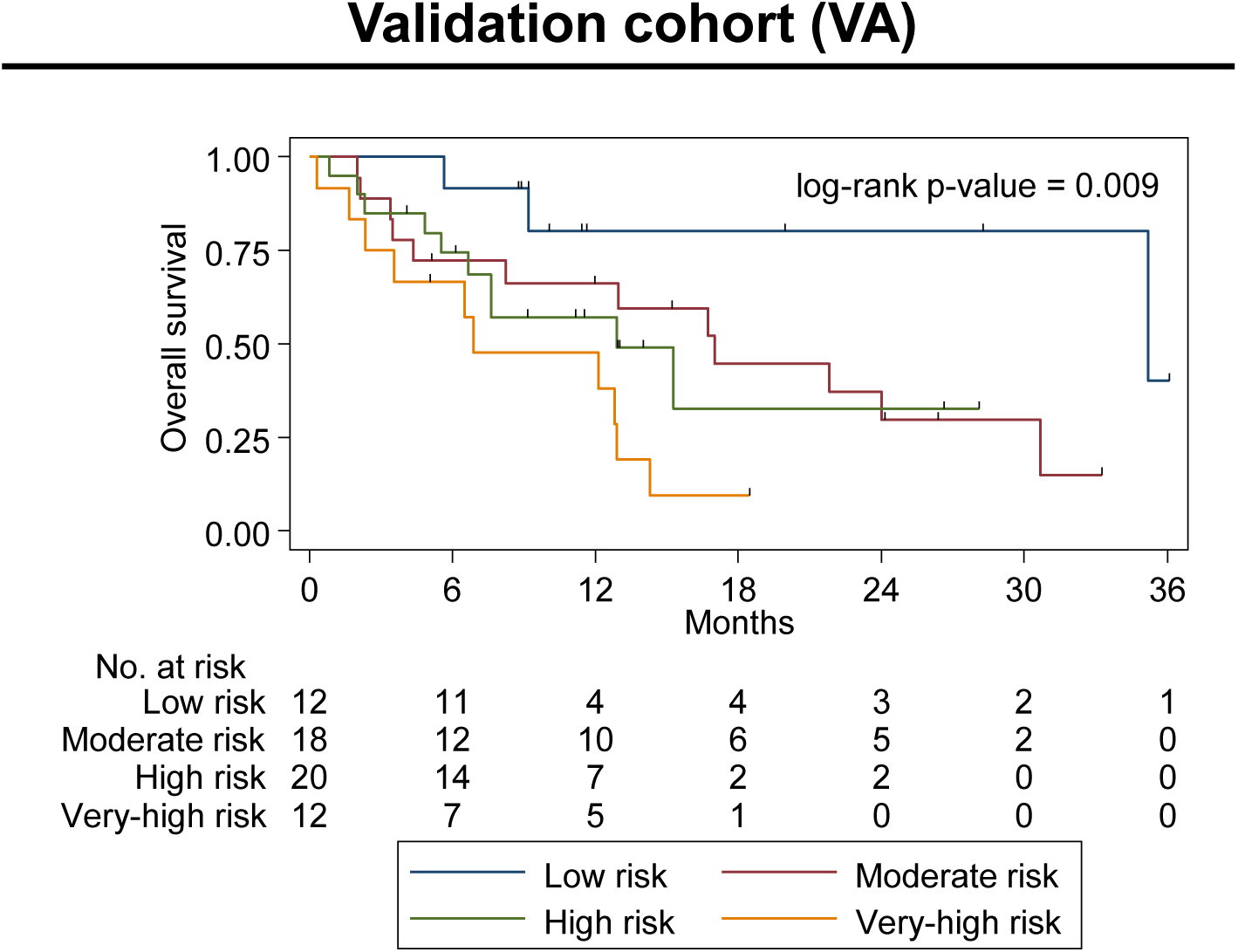
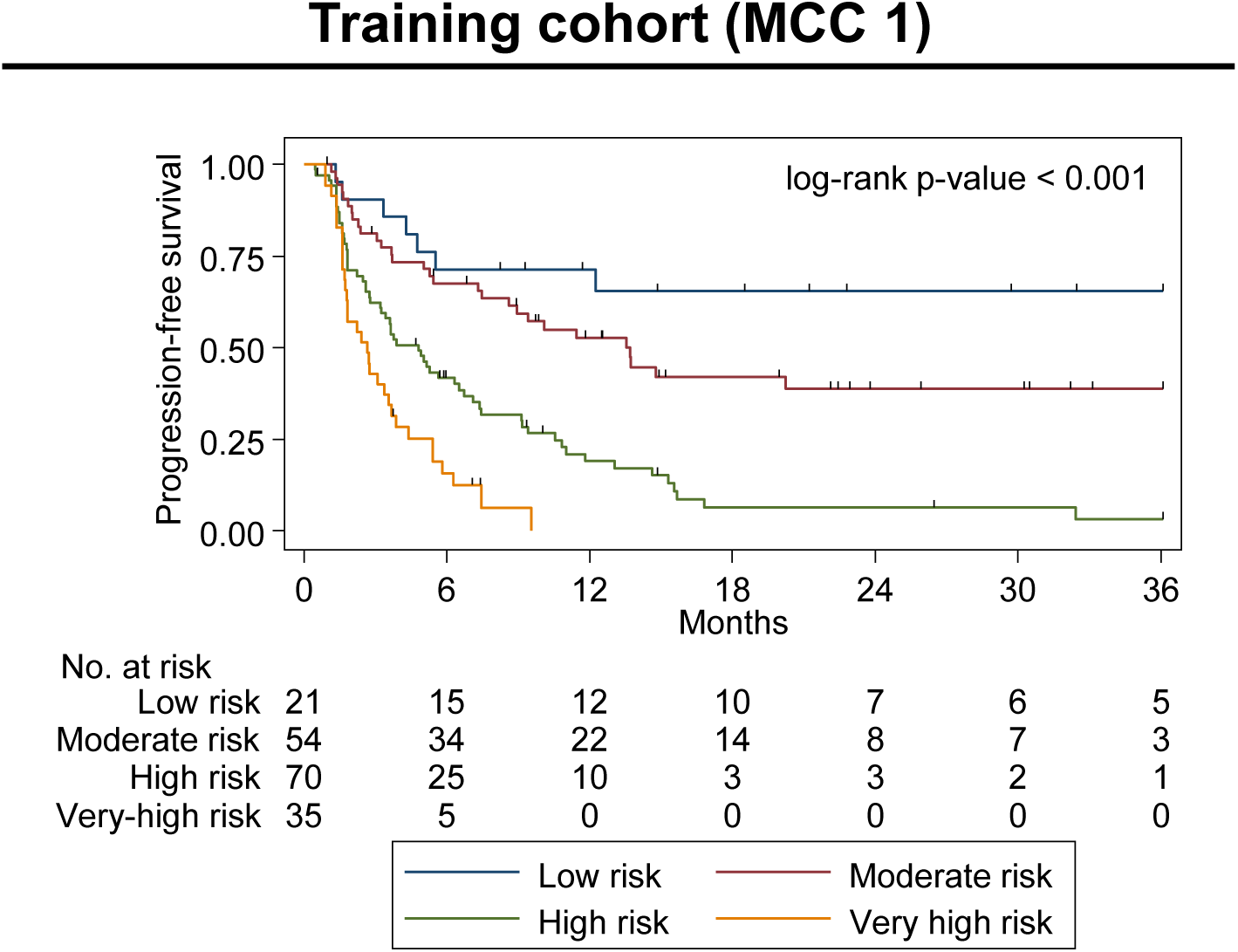
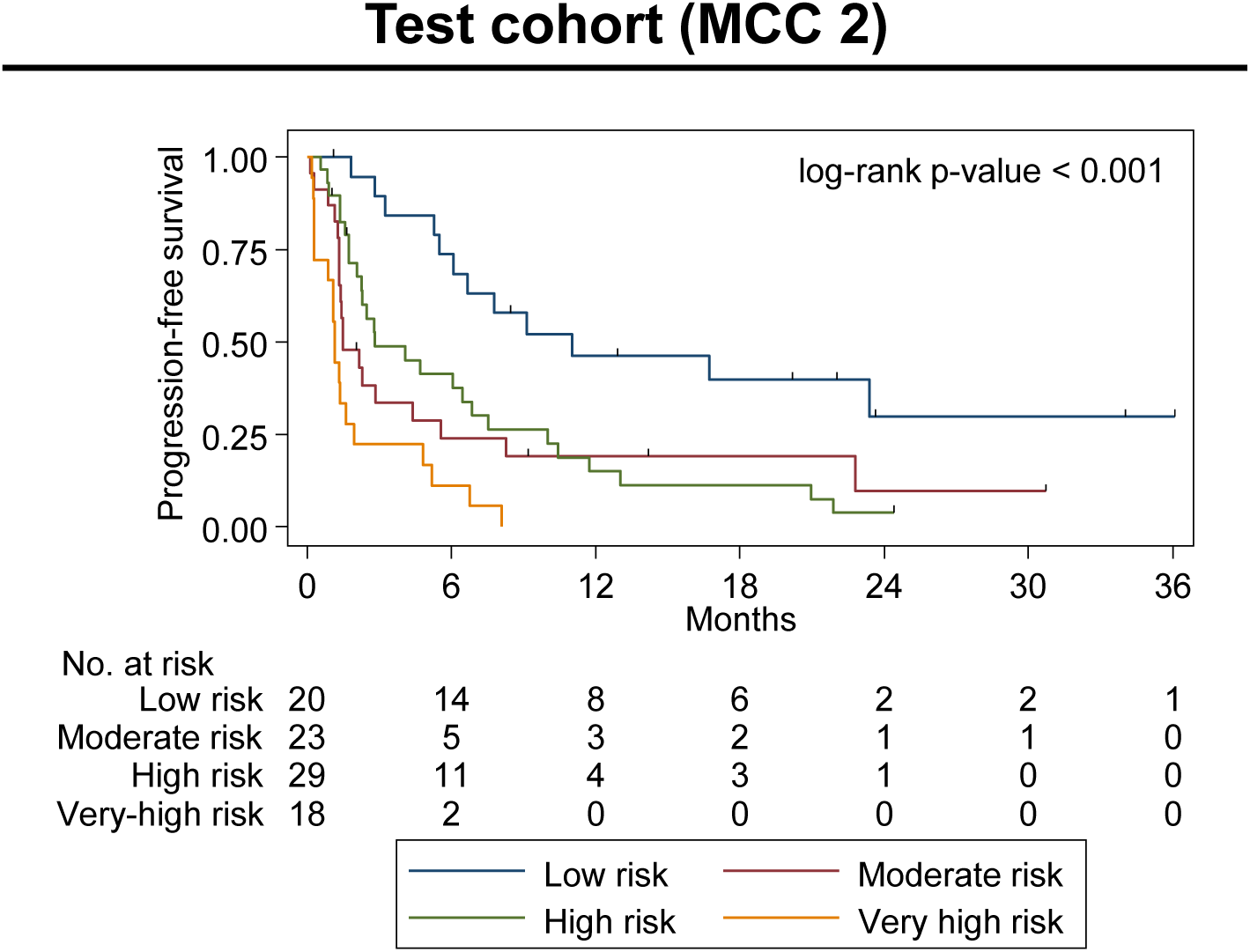
Kaplan-Meier survival curves estimates for overall survival between identified risk groups in the **A)** training (MCC 1) cohort, **B)** Test (MCC 2) cohorts and **C)** Validation (VA) cohort, and progressive-free survival in **D)** Training (MCC 1) cohort and **E)** Test (MCC 2) cohort. Test for agreement between radiomic and pathological immune response assessment.

### Multivariable analyses

Multivariable Cox regression analyses were conducted adjusting for clinical covariates that were significantly different between the training (MCC 1) and test (MCC 2) cohorts (Table 1). After adjusting for potential confounders, the high-risk (test cohort HR = 3.33; 95% CI 1.57 – 7.05) and very high-risk (test cohort HR = 5.35; 95% CI 2.14 – 13.36) groups were still found to be associated with significantly worse outcomes compared to the low-risk group (HR = 1.00). The results were consistent when the data were analyzed for PFS. Utilizing the validation cohort (VA) and adjusting for stage, ECOG, lymphocyte count and neutrophils to lymphocytes ratio; the very-high risk group had significantly worse outcomes (HR = 13.81; 95% CI 2.58 – 73.93) compared to the low-risk group (Supplementary Table 1).

Additionally, clinical covariates were compared across CART risk groups (Supplementary Table 2). Previous lines of therapy, ECOG, white blood cell counts, neutrophils and NLR were found to be significantly different between the CART groups (serum albumin and number of metastatic sites were not considered as potential confounders as these covariates were already part of the CART models). Multivariable Cox regression was performed adjusting for these potential confounders but did not appreciably alter the HRs for risk groups.

### Radiogenomics analyses

For two-group analysis, GLCM inverse difference was dichotomized at threshold determined by CART (0.43), which was found to be similar to the mean (0.47) and median (0.45) of GLCM inverse difference in the radiogenomics dataset. Correlation and two-group analyses identified 123 significant probesets representing 91 unique genes that were associated with the GLCM inverse difference radiomic feature (Supplemental Table 3). Pathway analysis indicated no significant enrichment (FDR < 0.05). Gene Ontology Biological Process enrichment of the gene set identified terms including regulation of cardiac conduction, sodium ion export across plasma membrane and membrane depolarization during action potential (Supplemental Table 4). Interestingly, only three probesets (representing two genes) were positively associated with GLCM inverse difference: carbonic anhydrase IX (CAIX) and Family with Sequence Similarity 83 Member F (FAM83F). GLCM inverse difference was positively associated with CAIX expression based on two different probesets (Fig. 4a-d). Median CAIX expression was lower for patients with low GLCM inverse difference (merck2-DQ892208_at: 4.61 [95% CI: 4.38 – 5.00]; merck-NM_001216_at: 4.48 [95% CI: 4.24 – 4.62]) vs. high GLCM inverse difference (merck2-DQ892208_at: 6.32 [95% CI: 5.50 – 6.86]; merck-NM_001216_at: 5.66 [95% CI: 5.11 – 6.39]).

**Figure 4.**
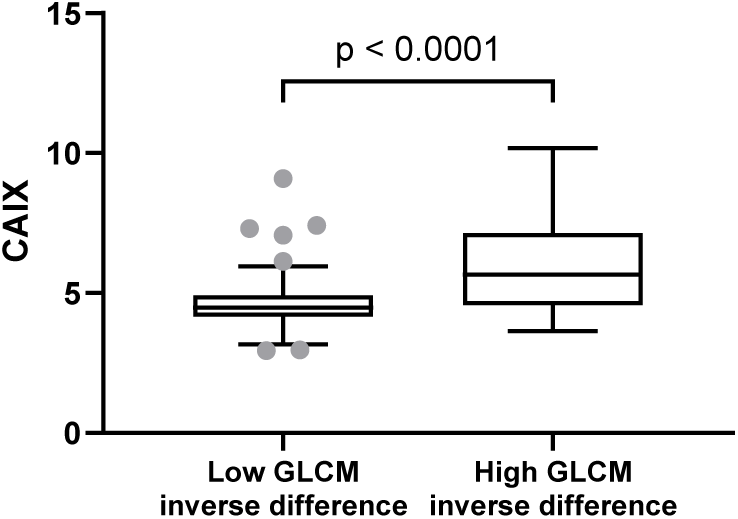
Association between GLCM inverse difference CT radiomic feature and CAIX expression. **A)** Whisker-box plots representing the association between CAIX expression using merck2-DQ892208_at probset and GLCM inverse difference. High and low GLCM inverse difference was found using novel cut-point (0.43) defined by CART analysis

**Figure 4B).**
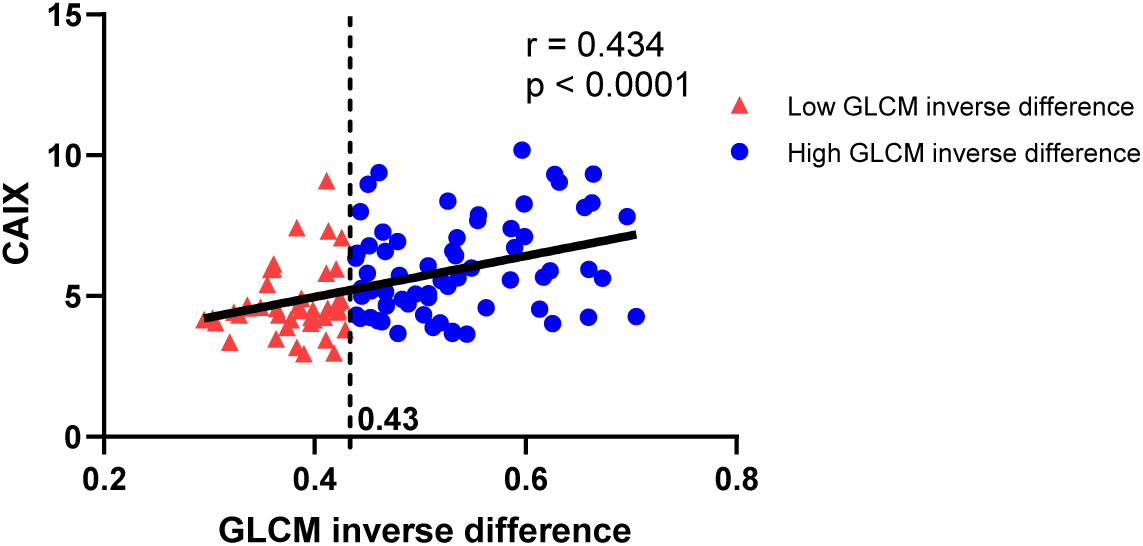
Scatter plot showing the linear relationship between CAIX expression using merck2-DQ892208_at probset and GLCM inverse difference. CART defined cut-off point was used to differentiate high (blue) and low (red) GLCM inverse difference.

**Figure 4C).**
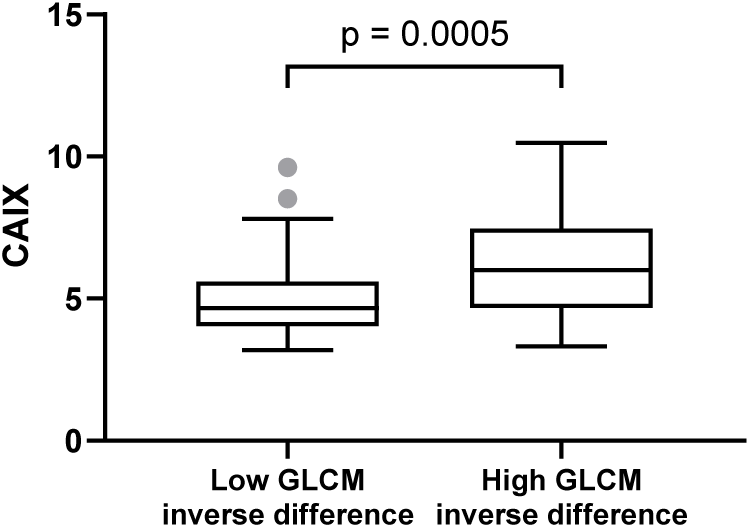
Whisker-box plots representing the association between CAIX expression using merck-NM_001216_at probset and GLCM inverse difference. High and low GLCM inverse difference was found using novel cut-point (0.43) defined by CART analysis.

**Figure 4D).**
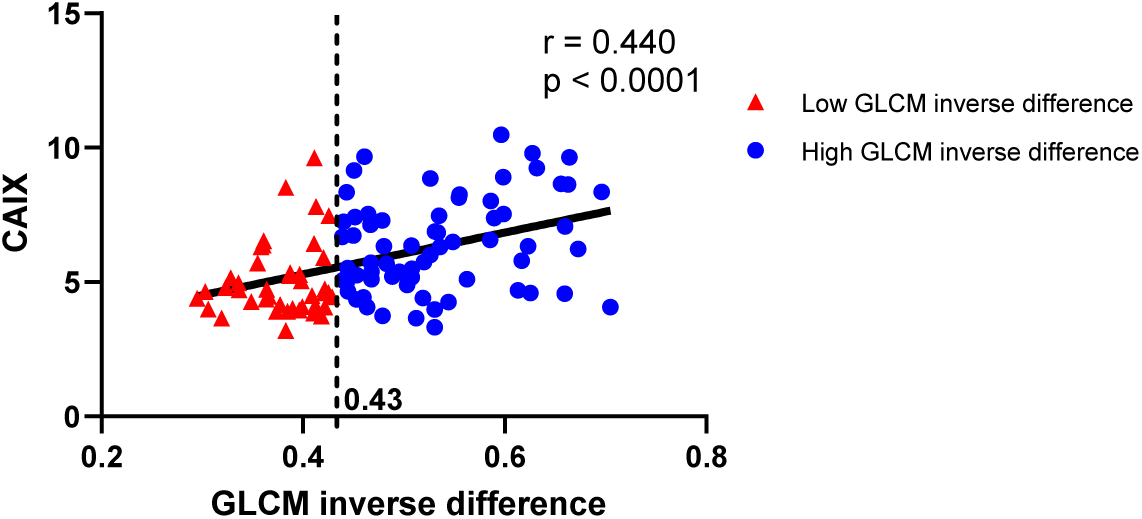
Scatter plot showing the linear relationship between CAIX expression using merck-NM_001216_at probset and GLCM inverse difference. CART defined cut-off point was used to differentiate high (blue) and low (red) GLCM inverse difference.

### Immunohistochemistry analyses

Analyses between CAIX IHC expression and GLCM inverse difference demonstrated that patients with higher CAIX expression had a borderline significant (p-value from Student’s *t*-test = 0.0646) higher GLCM inverse difference (Fig. 4e). The automated pathology scoring was highly correlated with the pathologist scored H-score (r = 0.6295, p = 0.0090, Fig. 4f). Representative cases of patients with high and low GLCM inverse difference and pathologically scored high and low IHC CAIX expressions are shown in Figure 4g.

**Figure 4E).**
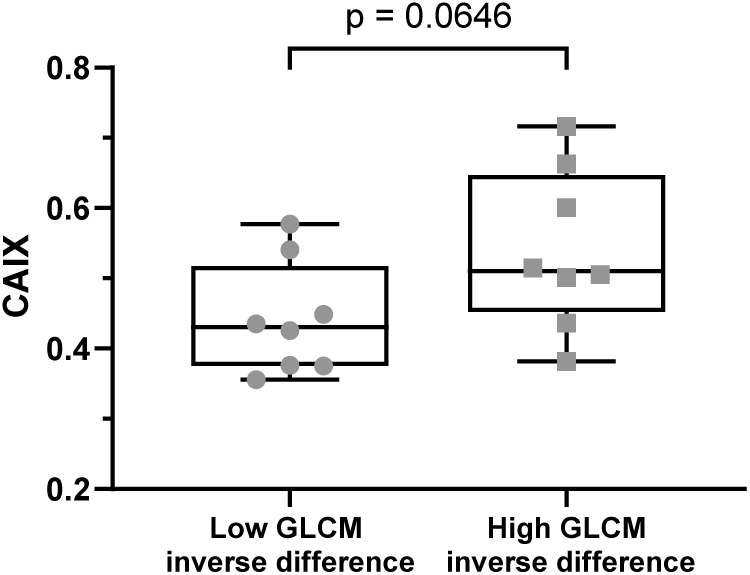
Whisker-box plots representing the association between CAIX expression on immunohistochemical staining and GLCM inverse difference CT radiomic feature. High and low GLCM inverse difference was found using novel cut-point (0.43) defined by CART analysis.

**Figure 4F).**
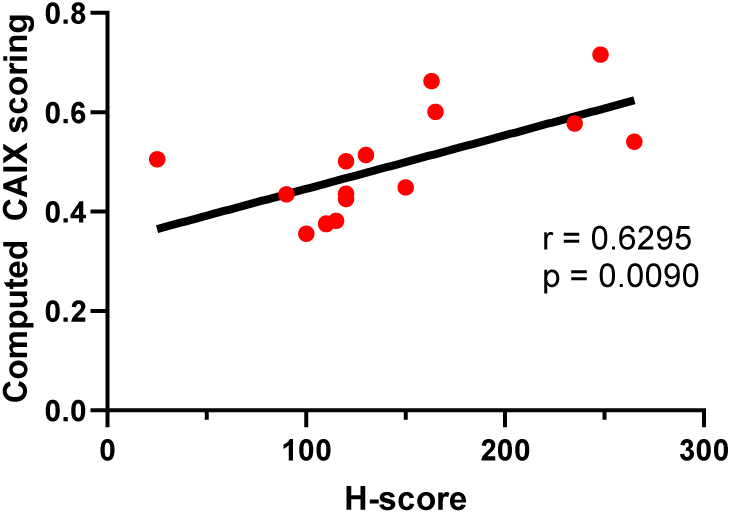
Scatter plot showing linear relationship between pathologist’s H-score for CAIX and computer derived (Aperio positive pixel count algorithm) automated CAIX scoring

**Figure 4G).**
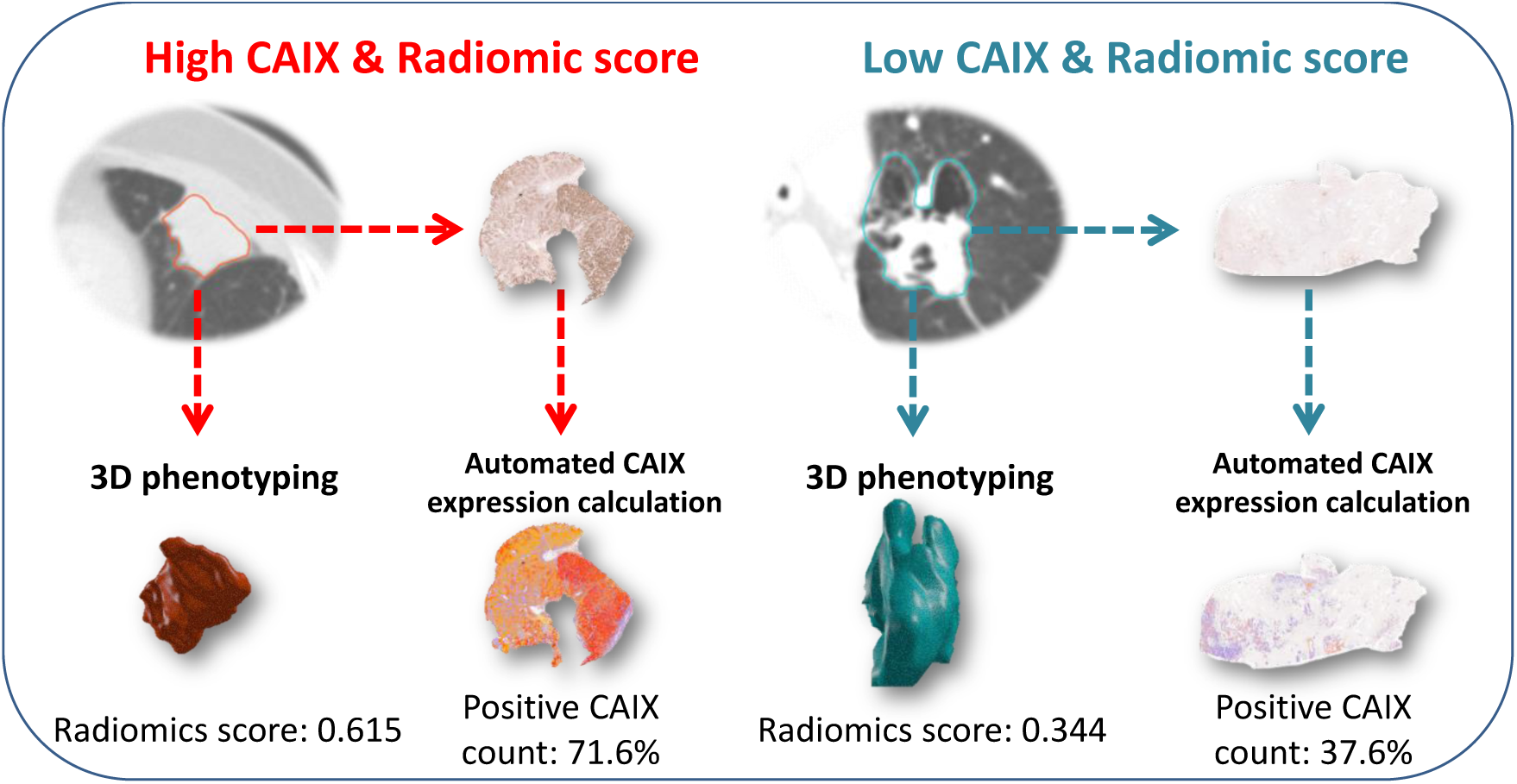
Representative cases for testing the agreement between GLCM inverse difference and CAIX IHC expression. Correlation between high CAIX and high CT radiomic feature is seen on left side and correlation between low CAIX and low CT radiomic feature is seen on right side.

### Prognostic validation datasets

GLCM inverse difference was significantly associated with OS in three out of the four independent NSCLC cohorts (Fig. 5a-g) using previously found CART cut-point (0.43). Although the *a priori* cut–point for GLCM inverse difference was not significantly associated with OS in the MAASTRO patient cohort, GLCM inverse difference as a continuous covariate was significantly associated with OS in a univariable Cox regression model (HR = 2.74; 95% CI 1.04 – 7.24).

**Figure 5.**
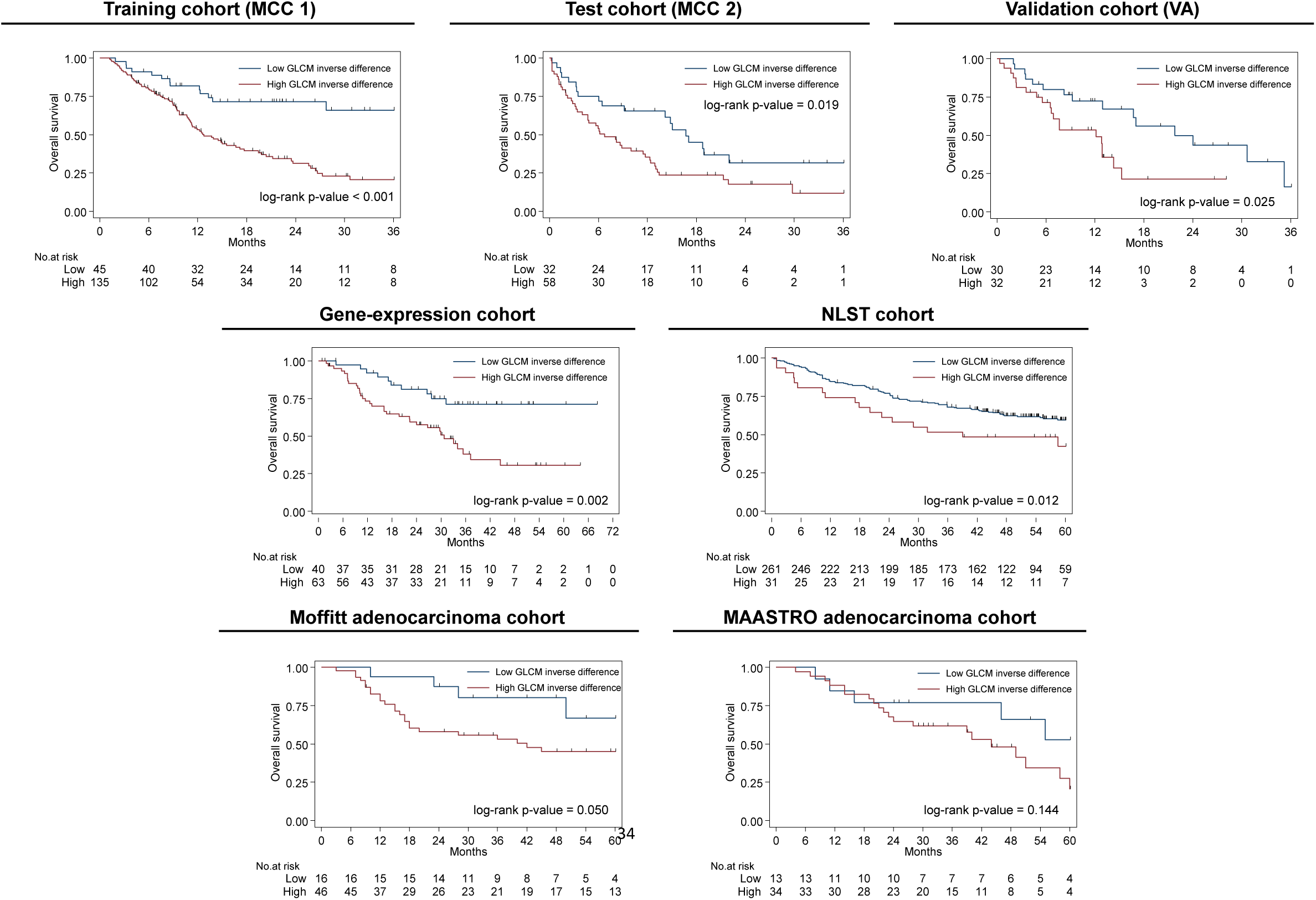
Kaplan-Meier survival plots of patients dichotomized by radiomics score. Same cut-off point was used for dichotomizing the cohorts. **A)** Training cohort (MCC 1) **B)** Test cohort (MCC 2) **C)** Validation cohort (VA) **D)** Gene-expression cohort **E)** Moffitt adenocarcinoma cohort **F)** MAASTRO adenocarcinoma cohort **G)** NLST cohort.

## Discussion

Predictive biomarkers that can identify lung cancer patients who are resistant to immunotherapy are critical unmet need as such, patients could avoid ineffective, unnecessary and expensive treatments. In this study, we utilized a rigorous radiomics pipeline and conducted robust analyses and identified a parsimonious clinical-radiomics model that significantly associates with survival outcomes. Patients were stratified into four unique risk groups (low, moderate, high and very-high risk) based on death and progression risk. The very high-risk group patients in the training cohort (MCC 1) had extremely poor OS which was tested and validated in independent test (MCC 2) and validation (VA) cohorts (Fig. 3). Using iRECIST criteria, patients in the very-high risk group were significantly more likely to develop progressive disease compared to patients in the low risk group (40.0% vs 4.8%). These findings suggest that patients in the very high-risk group should either avoid immunotherapy altogether and may be candidates for either chemotherapy or upfront combination treatments such as chemotherapy plus checkpoint blockade therapy. The most informative radiomic feature, GLCM inverse difference, was positively associated with CAIX expression and further validation demonstrated that GLCM inverse difference was also associated with OS in four independent NSCLC cohorts.

The four risk groups found in this study were derived from models using one radiomic feature (GLCM inverse difference) and two clinical covariates (baseline number of metastatic sites and serum albumin). As higher GLCM inverse difference alone was also associated with poor outcomes in four other prognostic NSCLC cohorts, it suggests a pan-radiomic feature. We characterized GLCM inverse difference as an “avatar” feature, representative of nine other highly correlated radiomic features (Fig. 2b). A higher GLCM inverse difference was associated with worse OS and it was observed in dense and uniform lesions (Supplementary Fig. 4). Assessment of the biological underpinnings revealed that this avatar feature was positively associated with CAIX expression on both gene-expression and IHC staining datasets. CAIX is an important pH regulatory enzyme that is upregulated in hypoxic tumors leading to an acidic and immunosuppressive tumor microenvironment^32^ and associate with poor prognosis^33,34^ including NSCLC^35,36^. Tumor-hypoxia leads to advanced but dysfunctional vascularization and acquisition of epithelial-mesenchymal transition phenotype, resulting in cell mobility and metastasis and alters cancer cell metabolism and contributes to therapy resistance by inducing cell quiescence and immunosuppressive phenotype^37^. The most predictive clinical covariates in this study demonstrate the utility of standard-of-care clinical information to predictive treatment response. Higher number of metastatic sites increases disease burden and can result in mixed responses where one or more lesions may be responding while others are progressing which ultimately results a progressive disease. The other clinical covariate, serum albumin, has shown to be associated with survival in NSCLC patients^38,39^ and used. As lower serum albumin is an indicator of malnutrition, inflammation, and hepatic dysfunction it may lead patients to a worse outcome. The mechanism of serum albumin with respect to immunotherapy response is yet to be established however, it was used in The Gustave Roussy Immune Score as a prognostic marker in immunotherapy phase I trials^40^. Emerging evidence demonstrates the utility of radiomics as a non-invasive approach to quantify and predict lung cancer treatment response of tyrosine kinase inhibitors^41,42^, platinum-based chemotherapy^43^, neo-adjuvant chemo-radiation^44,45^, stereotactic body radiation therapy^46,47^, and immunotherapy^8,48,49^. With respect to immunotherapy treatment response, our group previously demonstrated that pre-treatment clinical covariates and radiomic features predicted rapid disease progression phenotypes, including hyperprogression (AUROCs ranging 0.804-0.865) among 228 NSCLC patients treated with single agent or double agent immunotherapy^8^. Sun *et al.*^48^ developed and validated a radiomic signature for CD8 cells that predicted clinical outcomes (AUC = 0.67) among 135 patients treated with PD-1 or PD-L1 checkpoint blockades spanning 15 different cancer types. However, only 22% of their cohort was NSCLC patients. Trebeschi *et al.*^49^ developed a machine learning based model that discriminates progressive disease from stable disease and responsive disease (AUC = 0.83) between 123 NSCLC patients treated with PD-1 checkpoint blockade immunotherapy. As such, the study presented here represents the single largest study population of NSCLC patients treated with immunotherapy.

We do acknowledge some limitations of this study. TMB data were not available and PD-L1 IHC data were only available for 8 patients (4.4%) in the training cohort (MCC 1), 29 patients (32.2%) in the test cohort (MCC 2), and for no patients (0%) in the validation cohort (VA). Thus, we are unable to determine the performance of TMB or PD-L1 expression. However, recent studies have shown that, patients respond to immunotherapy regardless of PD-L1 expression^6,11^ or TMB, and both TMB and PD-L1 expression are prone to sampling artifact thus inclusion of PD-L1 expression or TMB may add little or no improvement to our prognostic models. Even though the training (MCC 1) and the test (MCC 2) cohorts were treated at the same hospital, there were significant OS differences as majority of patients in the test cohort (MCC 2) were treated with standard-of-care immunotherapy while the training cohort (MCC 1) comprised of clinical trial patients who typically have higher performance status (Supplementary Fig. 1). However, patients sorted in the very-high risk group consistently had worse survival outcomes in both test (MCC 1) and validation (VA) cohorts. Finally, potentially important covariate for immune response, lactic acid dehydrogenase (LDH) data, was only available for a subset of patients in the training (N=162) and test cohorts (N = 14) hence, we did not include LDH in our analyses. However, when the risk group HRs were adjusted for LDH, the HR for the very-high risk group (HR = 15.38, 95% CI: 4.57 – 51.83) did not change significantly and LDH was not significantly associated with OS (multivariable Cox regression p-value = 0.992). Despite these minor weaknesses, this study yields a high radiomic quality score (RQS = 22)^31^ (Supplementary Table 5) which is a stringent metric that quantifies the clinical relevance of a radiomic study.

In conclusion, using standard-of-care imaging and clinical covariates we identified and validated a novel parsimonious model that is associated with OS and PFS among NSCLC patients treated with immunotherapy. The prognostic image-based (i.e., GLCM inverse difference) feature was found to be related to CAIX, an important enzyme upregulated in hypoxic and acidotic tumors which is linked to treatment resistance. The potential clinical application of this work is to identify patients that are resistant to immunotherapy to avoid ineffective, unnecessary and expensive treatments using readily available clinical covariates and medical images. Future studies are needed to find potential relevance of these models in other cancer sites.

## Acknowledgments

Funding support came from the National Cancer Institute (NCI) (U01-CA143062 to Drs. Gillies and Schabath), the NCI Early Detection Research Network (U01-CA200464 to Drs. Gillies and Schabath), and CA186145 and CA196405. This work has also been supported in part by a Cancer Center Support Grant (CCSG) at the H. Lee Moffitt Cancer Center and Research Institute; an NCI designated Comprehensive Cancer Center [grant number P30-CA76292]. Also, this material is the result of work supported with resources and the use of facilities at the Moffitt Cancer Center and the James A. Haley Veterans’ Hospital.

We also would like to thank to the Image Response Assessment Team (IRAT) Shared Resource Core for their assistance in implementing the IBSI radiomic feature set.

## Code availability statement

The MATLAB^®^ scripts to create peritumoral masks from intratumoral masks are publically available at https://github.com/TunaliIlke/peritumoral_regions/. The radiomic features were extracted using algorithms from the Image Biomarker Standardization Initiative (IBSI) v5^26^. The algorithms were written in C++ and MATLAB^®^ v2015a, and these codes are available upon request from the corresponding author (M.B.S.).

## Data availability statement

The data and data analysis code will be available upon reasonable request from the corresponding author (M.B.S.).

## Disclosure of Potential Conflicts of Interest

J.E. Gray reports receiving commercial research grants from AstraZeneca, Merck, Array, Epic Sciences, Genentech, Bristol-Myers Squibb, BI, Trovagene, and Novartis, and is a consultant/advisory board member for AstraZeneca, Janssen, Genentech, Eli Lilly, Celgene, and Takeda, and other remuneration from Genentech, AstraZeneca, Merck, and Lilly/Genenech. R.J. Gillies is an investor and member of the Advisory Board at HealthMyne, Inc., and has Research support form Helix Biopharma. No potential conflicts of interest were disclosed by the other authors.

Contents of this research do not represent the views of the Department of Veterans Affairs or the United States Government.

