## Supplemental Methods for "Hypoxia-related radiomics predict immunotherapy response: A multi-cohort study of NSCLC"

***Immunotherapy-treated lung cancer patient population and data***

Common inclusion criteria for **clinical trial** patients included: patients who were diagnosed with histologically- or cytological-documented NSCLC with advanced/metastatic stage disease with at least one measurable lesion (≥ 10 mm), Eastern Cooperative Oncology Group (ECOG) Performance Status of 0 or 1, and provided written informed consent. Common exclusion criteria included: a concurrent medical condition requiring the use of immunosuppressive medications or immunosuppressive doses of systemic or absorbable topical corticosteroids, and presence of any active autoimmune disease. Access to these data for retrospective analyses was approved by the University of South Florida Institutional Review Board.

Moffitt’s Cancer Registry abstracts information from patient electronic medical records on demographics, history of smoking, stage, histology, treatment, and vital status. Follow-up for vital status occurs annually through active (i.e., chart review and directly contacting the patient, relatives, and other medical providers) and passive methods (i.e., mortality records). Hematology data were obtained from the CDCS and included: lactate dehydrogenase (LDH), serum albumin, lymphocytes, white blood cells (WBC), neutrophils, fibrinogen, and neutrophil to lymphocyte ratio (NLR). Manually abstracted data included: targeted mutations (*EGFR, KRAS*), history of systemic treatment(s) for current lung cancer staging, ECOG performance status, number of metastatic sites (number of organs that have metastatic lesions), and metastatic sites prior to treatment.

All patient data collected from James A. Haley Veterans’ Hospital (VA cohort) were manually collected from electronic medical records of the patients.

**Selection of stable and reproducible features**

Two separate publicly available datasets (downloaded from: <http://www.cancerimagingarchive.net>) were utilized to assess stability (The Moist-run dataset ^1^), and reproducibility (RIDER dataset ^2^) of radiomic features to increase the likelihood of a reproducible and robust radiomics model.

The Moist-run dataset was constructed by the Quantitative Imaging Network (QIN) as part of a lung segmentation challenge ^1^ and consists of 40 chest CT images of 40 NSCLC patients and one thoracic phantom from five collections of Digital Imaging and Communications Medicine series. Each patient in the dataset had one lesion of interest and the thoracic phantom scan had 12 lesions of interest. The RIDER test-retest dataset which was used to find the reproducible features ^2^ consisted of 32 NSCLC patients with two separate non-contrast CT scans acquired within 15 minutes of each other using the same scanner with fixed acquisition and processing parameters. As such, the only variation between the test and retest scans were attributed to patient orientation, respiratory, and movement. The images on these datasets were previously de-identified

Using the Moist run dataset, all radiomic features were computed for 9 different segmentations done by 3 different algorithms which each were run by 3 different initial parameters. Afterwards, concordance correlation coefficient (CCC) metric was calculated to assess inter- and intra-segmentation differences of the radiomic features. The RIDER dataset was utilized to assess reproducibility of radiomic features between test and re-test scans. After extracting radiomic features from both scans of the patients, CCC values were calculated and features that have a CCC < 0.75 were eliminated. Shape features were only extracted from intratumoral regions as they were proven to be highly correlated (Pearson correlation > 0.95) with their peritumoral versions. Additionally details on stable and reproducible radiomic feature selection is described elsewhere^3^.

**Staining procedure for CAIX (human expression)**

Immunohistochemical staining:  Slides were stained using a Ventana Discovery XT automated system (Ventana Medical Systems, Tucson, AZ) as per manufacturer's protocol with proprietary reagents. Briefly, slides were deparaffinized on the automated system with EZ Prep solution (Ventana). Heat-induced antigen retrieval method was used in RiboCC (Ventana). The rabbit primary antibody that reacts to CA-IX, (#ab15086, Abcam, Cambridge, MA) was used at a 1:250 concentration in Dako antibody diluent (Carpenteria, CA) and incubated for 32 min. The Ventana OmniMap Anti-Rabbit Secondary Antibody was used for 20 min. The detection system used was the Ventana ChromoMap kit and slides were then counterstained with Hematoxylin.  Slides were then dehydrated and cover slipped as per normal laboratory protocol.
