## Supplemental Tables 1-2 for "Hypoxia-related radiomics predict immunotherapy response: A multi-cohort study of NSCLC"

| **Sup Table 1**. Univariable and multivariable Cox regression analysis for overall survival for the validation cohort. | | | | |
| --- | --- | --- | --- | --- |
|  | |  | **Validation cohort (VA)**  **(N = 62)** | |
| **Overall survival** | |  | **Univariable Model^1^**  **HR (95% CI)** | **Multivariable Model^2^**  **HR (95% CI)** |
| **Risk group** | |  |  |  |
|  | Low-risk |  | *1.00 (Reference)* | *1.00 (Reference)* |
|  | Moderate-risk |  | 4.07 (0.90 – 18.26) | 4.00 (0.83 – 19.17) |
|  | High-risk |  | **4.72 (1.02 – 21.94)** | 4.54 (0.90 – 23.11) |
|  | Very-high-risk |  | **9.72 (2.08 – 45.49)** | **13.81 (2.58 – 73.93)** |
| **Age** | |  | . | 1.00 (0.96 – 1.05) |
| **Stage** | |  | . | 0.66 (0.23 – 1.93) |
| **ECOG PS** | |  | . | 1.98 (0.99 – 3.94) |
| **Lymphocytes** | |  | . | 1.13 (0.53 – 2.40) |
| **NLR** | |  | . | 1.09 (0.99 – 1.18) |
| Abbreviations: SD = standard deviation; HR = hazard ratio; CI = confidence interval; NLR = neutrophils to lymphocytes ratio;  **Bold** values are statistically significant.  ^1^The main effects for each risk group with the low-risk group as the referent category.  ^2^The low-risk group was used as the referent group and the models were adjusted for clinical covariates that were found to be significantly different between the training and validation cohorts (Table 1). | | | | |

| **Sup. Table 2.** Patient characteristics by CART risk groups for the training cohort. | | | | | | |
| --- | --- | --- | --- | --- | --- | --- |
| **Characteristic** | | **Low-risk** | **Moderate-risk** | **High-risk** | **Very-high-risk** | **P-Value** |
| **Age at diagnosis, N (%)** | |  |  |  |  |  |
|  | ***Dichotomized*** |  |  |  |  |  |
|  | < 65 | 9 (42.9) | 22 (40.7) | 26 (37.1) | 11 (31.4) |  |
|  | ≥ 65 | 12 (57.1) | 32 (59.3) | 44 (62.9) | 24 (68.6) | 0.798 |
| **Sex, N (%)** | | | |  |  |  |
|  | Female | 8 (38.1) | 31 (57.4) | 40 (57.1) | 16 (45.7) |  |
|  | Male | 13 (61.9) | 23 (42.6) | 30 (42.9) | 19 (54.3) | 0.323 |
| **Smoking status, N (%)** | |  |  |  |  |  |
|  | Never smoker | 4 (19.1) | 10 (18.9) | 12 (17.9) | 4 (11.4) |  |
|  | Ever smoker | 17 (80.9) | 43 (81.1) | 55 (82.1) | 31 (88.6) | 0.809 |
| **Stage, N (%)** | | | |  |  |  |
|  | III | 2 (9.5) | 0 (0) | 3 (4.3) | 1 (2.9) |  |
|  | IV | 19 (90.5) | 54 (100) | 67 (95.7) | 34 (97.1) | 0.138 |
| **Histology, N (%)** | |  |  |  |  |  |
|  | Adenocarcinoma/others | 17 (81.0) | 43 (79.6) | 53 (75.7) | 24 (68.6) |  |
|  | Squamous cell carcinoma | 4 (19.0) | 11 (20.4) | 17 (24.3) | 11 (31.4) | 0.636 |
| **Checkpoint inhibitors, N (%)** | |  |  |  |  |  |
|  | Anti PD-L1 | 4 (19.1) | 11 (20.37) | 24 (34.3) | 9 (25.7) |  |
|  | Anti PD-1 | 7 (33.3) | 16 (29.6) | 25 (35.7) | 9 (25.7) |  |
|  | Doublet | 10 (47.6) | 27 (50.0) | 21 (30.0) | 17 (48.6) | 0.285 |
| **ECOG performance status, N (%)** | | | |  |  |  |
|  | 0 | 10 (47.6) | 10 (18.5) | 15 (21.4) | 4 (11.4) |  |
|  | 1 | 11 (52.4) | 44 (81.5) | 55 (78.6) | 31 (88.6) | **0.021** |
| **Previous lines of therapy on current diagnosis, N (%)** | | | |  |  |  |
|  | None | 10 (47.6) | 33 (61.1) | 10 (14.3) | 17 (48.6) |  |
|  | 1 | 4 (19.1) | 13 (24.1) | 24 (34.3) | 7 (20.0) |  |
|  | ≥ 2 | 7 (33.3) | 8 (14.8) | 36 (51.4) | 11 (31.4) | **<0.001** |
| **Number of metastatic sites^1^, N (%)** | |  |  |  |  |  |
|  | 1 | 21 (100) | 14 (25.9) | 47 (67.1) | 0 (0) |  |
|  | ≥ 2 | 0 (0) | 40 (74.1) | 23 (32.9) | 35 (100) | **<0.001** |
| ***EGFR* mutational status, N (%)** | |  |  |  |  |  |
|  | Not Detected | 14 (77.8) | 36 (87.8) | 37 (75.5) | 20 (83.3) |  |
|  | Detected | 4 (22.2) | 5 (12.2) | 12 (24.5) | 4 (16.7) | 0.495 |
| ***KRAS* mutational status, N (%)** | |  |  |  |  |  |
|  | Not Detected | 7 (58.3) | 17 (60.7) | 26 (70.3) | 11 (84.6) |  |
|  | Detected | 5 (41.7) | 11 (39.3) | 11 (29.7) | 2 (15.4) | 0.401 |
| **Hematology,** **median, (95% CI)** | |  |  |  |  |  |
|  | Serum albumin^1^, (g/dL) | 4.0 (3.8-4.2) | 4.1 (4.1-4.2) | 4.0 (3.9-4.1) | 3.6 (3.5-3.7) | **<0.001** |
|  | Lymphocytes, (1e+9/L) | 1.0 (0.8-1.4) | 1.2 (1.2-1.4) | 1.4 (1.3-1.5) | 1.2 (0.8-1.6) | 0.215 |
|  | WBC, (1e+9/L) | 6.8 (5.1-8.8) | 6.9 (6.4-8.0) | 6.9 (6.4-7.4) | 8.3 (7.4-10.9) | **0.023** |
|  | Neutrophils, (1e+9/L) | 4.8 (3.7-6.4) | 4.7 (4.1-5.3) | 4.4 (3.9-4.9) | 6.1 (5.1-7.4) | **0.007** |
|  | Ratio of: Neutrophils/Lymphocytes | 4.1 (2.7-5.7) | 3.4 (2.8-4.0) | 3.1 (2.8-3.7) | 4.6 (3.8-7.0) | **0.004** |
| Abbreviations: CI = confidence interval;  **Bold** P-values are statistically significant and P-values for continuous variables were calculated using Kruskall-Wallis test and Fisher’s Exact Test of categorical variables.  ^1^Serum albumin and number of metastatic sites were already part of the CART models. | | | | | | |
