## Supplemental Figures for "Hypoxia-related radiomics predict immunotherapy response: A multi-cohort study of NSCLC"

**Supplementary Figure Legends**

**Sup Figure 1**.	**Overall survival and progression-free survival Kaplan-Meier graphs for the training (MCC 1), test (MCC 2) and validation (VA) cohorts.** Progression-free survival data for the validation cohort (VA) was not available.

**Sup Figure 2**. **Overall survival and progression-free survival Kaplan-Meier graphs for initial risk groups identified by CART in the training cohort.** Groups 2 and 3, and groups 4 and 5 were later combined (Figure 3a and d).

**Sup Figure 3**. **Time-dependent AUC curves for Cox regression models based on 6, 12, 24 and 36 months for training (top), test (middle) and validation cohorts (bottom).** The AUC values were not statistically different between training and test cohorts.

**Sup Figure 4**. **Two patients marked as low (top) and very-high (bottom) risk group.** First column represents the primary target lesion CT scan. Second column represents the tumor segmentation. Third column represents a radial gradient image of the segmented area for visualization of tumor texture. Patient marked as a low-risk had a less dense tumor phenotype with a lower GLCM inverse difference score. Patient marked as very-high risk had a dense tumor phenotype with a higher GLCM inverse score.

Sup Figure 1

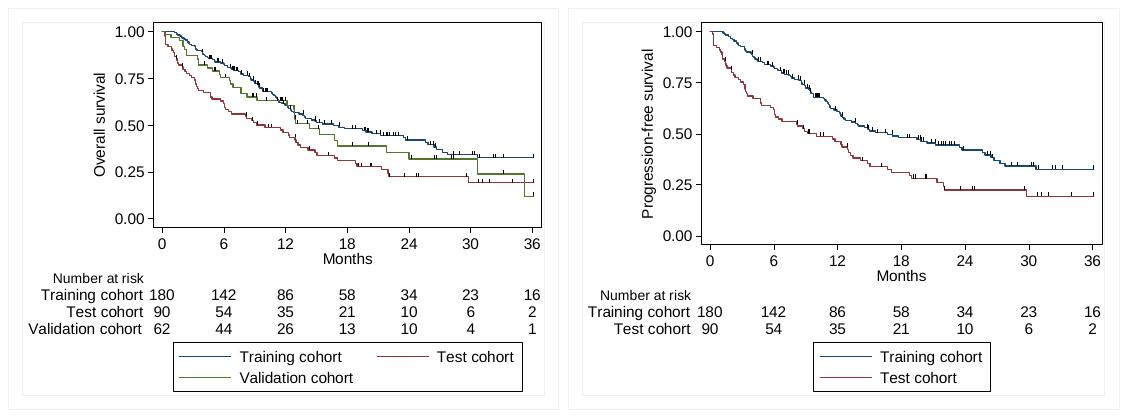

Sup Figure 2

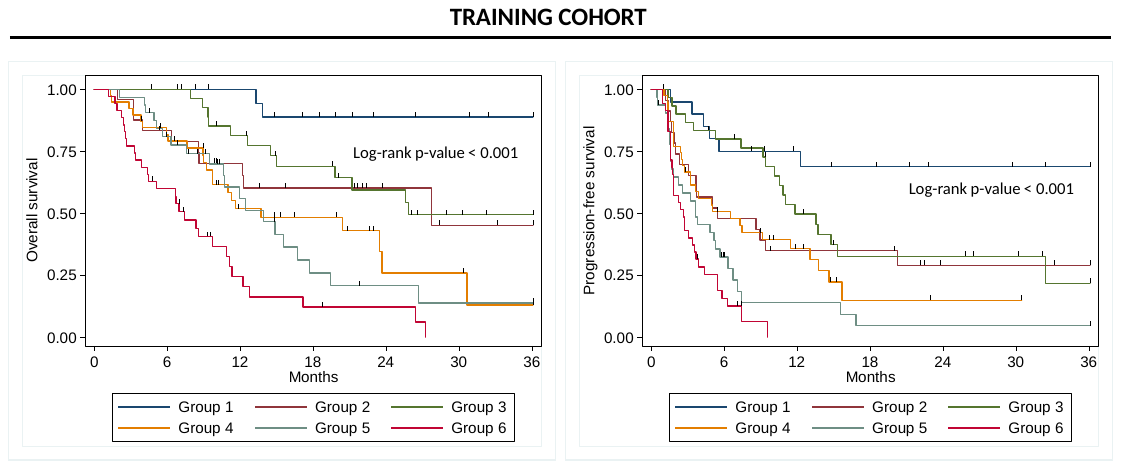

Sup Figure 3

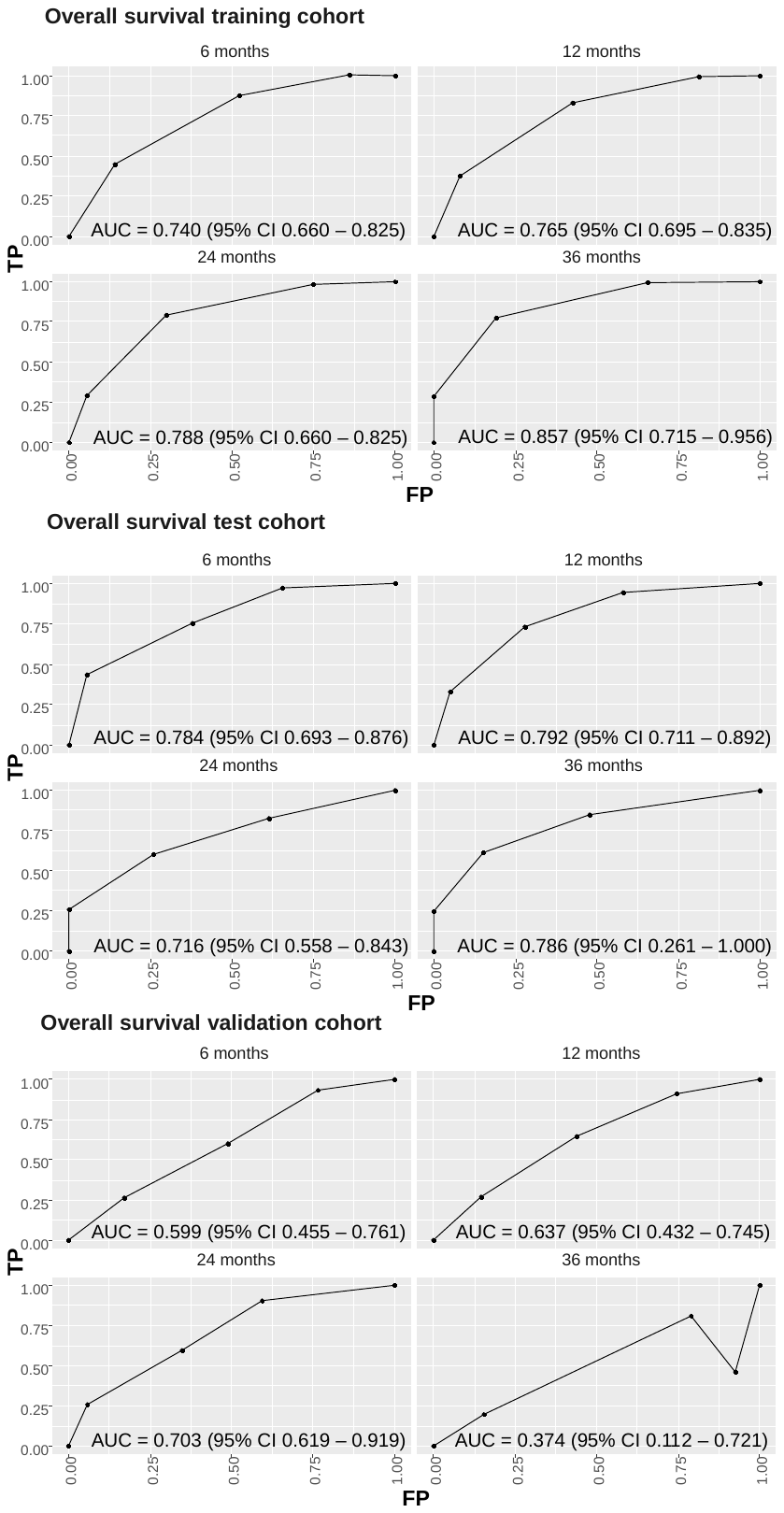

Sup Fig 4

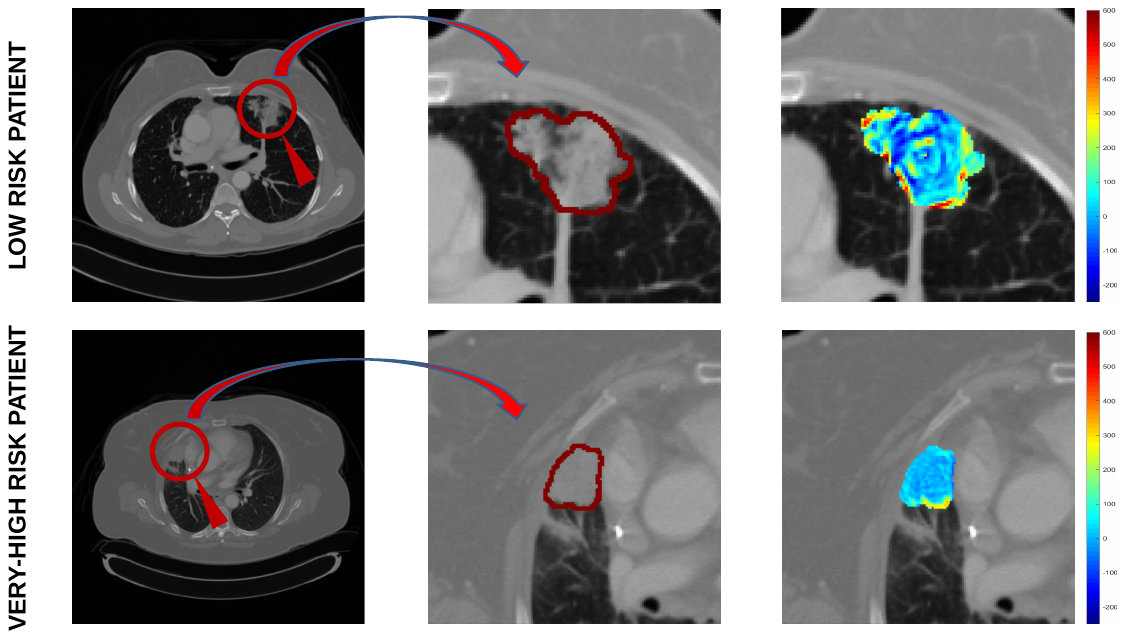
